# Data-driven Identification of Total RNA Expression Genes (TREGs) for Estimation of RNA Abundance in Heterogeneous Cell Types

**DOI:** 10.1101/2022.04.28.489923

**Authors:** Louise A. Huuki-Myers, Kelsey D. Montgomery, Sang Ho Kwon, Stephanie C. Page, Stephanie C. Hicks, Kristen R. Maynard, Leonardo Collado-Torres

## Abstract

Next-generation sequencing technologies have facilitated data-driven identification of gene sets with different features including genes with stable expression, cell-type specific expression, or spatially variable expression. Here, we aimed to define and identify a new class of “control” genes called Total RNA Expression Genes (TREGs), which correlate with total RNA abundance in heterogeneous cell types of different sizes and transcriptional activity. We provide a data-driven method to identify TREGs from single cell RNA-sequencing (RNA-seq) data, available as an R/Bioconductor package at https://bioconductor.org/packages/TREG. We demonstrated the utility of our method in the postmortem human brain using multiplex single molecule fluorescent in situ hybridization (smFISH) and compared candidate TREGs against classic housekeeping genes. We identified *AKT3* as a top TREG across five brain regions, especially in the dorsolateral prefrontal cortex.

## Background

In genomic analyses, researchers frequently face the decision on whether to use a list of genes identified *a priori* for an analysis or to identify new genes in a data-driven manner that have specific desirable qualities to answer a biological question. This duality reflects the nature of how our knowledge evolves as experimental assays generate more data and provide further insight into our understanding of biological systems. This expansion of knowledge is reflected in approaches such as single cell or nucleus RNA sequencing (sc/snRNA-seq) where known cell type marker genes are used to annotate cells, and the annotations are used to find new cell type marker genes [1–3]. Similarly, in spatially-resolved transcriptomics, previous knowledge of genes with distinct spatial expression can be used to annotate cells *in situ*, but also identify anatomical domains leading to identification of new spatially variable gene sets [4, 5].

Methods for gene selection, either data-driven or based on previous knowledge from the literature [6], are not only relevant to genes with high variability, but also to identify “control” genes with stable levels of expression used, for example, in normalization, such as microarray channel [7] or quantitative PCR normalization [8]. One data-driven approach to identify control genes for these assays when samples contain different amounts of RNA is to rely on a rank-invariant approach [9].

Different cell types contain variable amounts of RNA due to differences in cell size and transcriptional activity. In brain tissue, this variation in cell size and RNA abundance can negatively impact the accuracy of bulk RNA-seq deconvolution methods, which aim to identify cell type proportions in homogenate tissue by using sc/snRNA-seq reference profiles [10]. For example, neurons are larger and more transcriptionally active than glia and therefore have more RNA content and more genes detected per nucleus in snRNA-seq data [11]. With the exception of one method [12], the majority of existing bulk RNA-seq deconvolution methods [10] fail to incorporate this variation and hence report potentially biased estimates of the *relative* fraction of RNA attributable to each cell type rather than the true proportion of cell types [13]. However, methods to robustly estimate cell or nuclear size and total RNA abundance in the same assay are limited as approaches that capture global RNA expression, such as snRNA-seq, require tissue homogenization preventing the acquisition of cell size measurements.

One approach to measure nuclear size and relative RNA abundance in the same assay is to use multiplex single-molecule fluorescent *in situ* hybridization (smFISH) using the RNAscope technology [14], which allows quantification of both cell morphology and gene expression for a small number of target genes. Specifically, RNAscope fluorescently labels individual RNA transcripts, which are represented as ‘dots’ or puncta in the image that can be segmented and used to quantify gene expression per nucleus [15]. In parallel, these images can be used to estimate spatially-resolved nuclear size across heterogenous cell types *in situ*. However, there are no rigorous and data-driven approaches to identify candidate target genes to estimate total RNA abundance compatible with smFISH.

Here, we propose a data-driven approach using sc/snRNA-seq data to identify a class of genes we refer to as Total RNA Expression Genes (TREGs) to estimate total RNA abundance in heterogeneous cell types. These genes should ideally be highly correlated with total RNA abundance and predictive of transcriptional activity (**Figure 1a**). In the postmortem human brain, single unit measurements are limited to the nucleus, but it has been established that nuclear RNA content is representative of the whole cell [16]. In other research settings, single unit measurements could encompass the whole cell using scRNA-seq. When TREGs are applied in smFISH using RNAscope, they can be used to link spatially-resolved size and total RNA expression in different cells.

**Figure 1:**
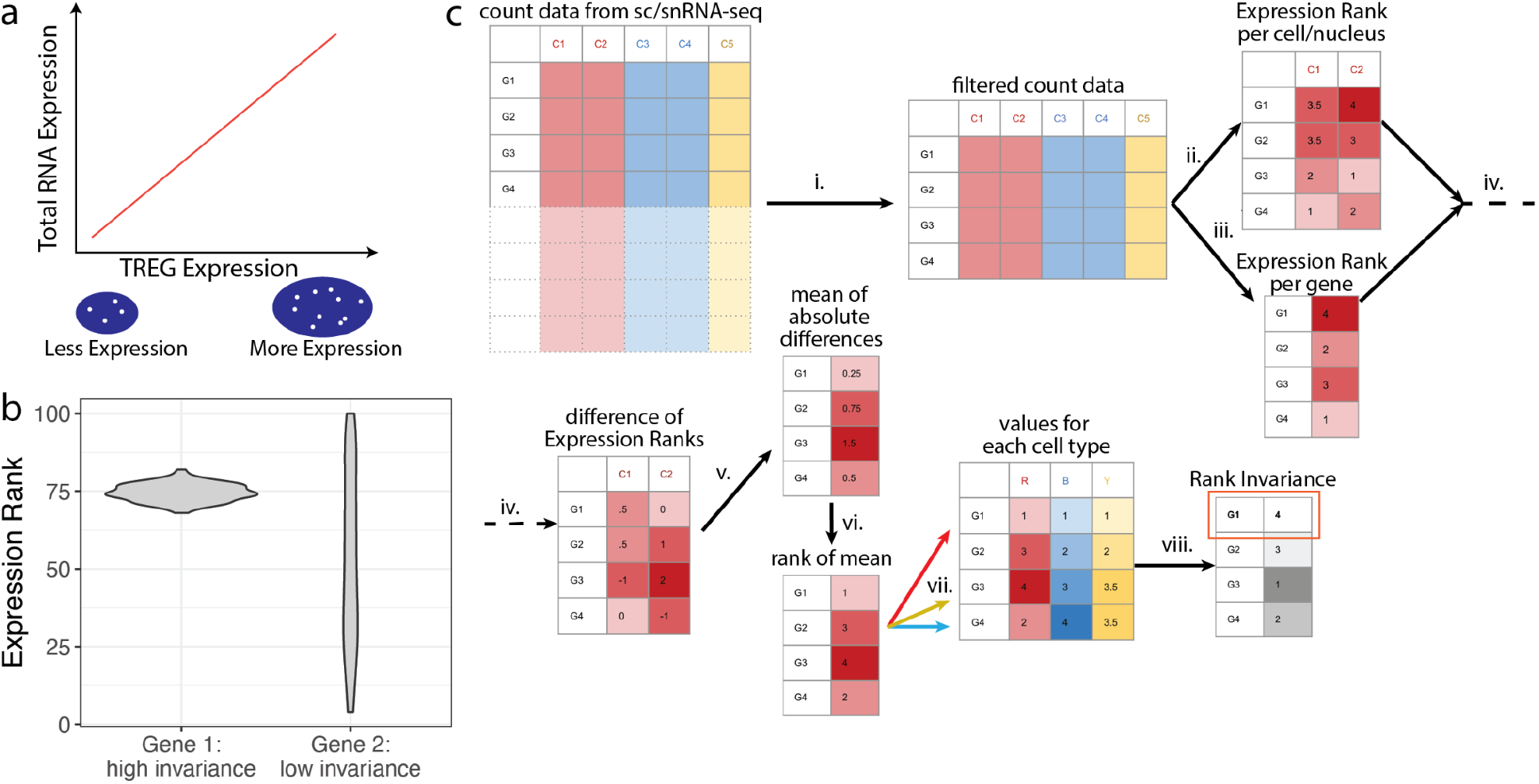
Overview of TREG motivation and methodology. **a.** Illustration of the relationship between the expression of a TREG and the total RNA expression of a nucleus. TREG expression can be quantified with puncta (white dots) in a nucleus (blue area), where the nucleus is identified with DAPI. **b.** Illustration of the distribution of Expression Rank, which is the rank of the expression of a given gene among all genes, computed individually for each cell/nucleus, depending on the measurement technology used: sc or snRNA-seq. Two theoretical genes are shown, Gene 1 with high Rank Invariance and Gene 2 with low Rank Invariance across cells/nuclei. **c.** Rank Invariance workflow to identify a TREG (Methods: Rank invariance calculation), with a gene expression matrix with genes on the rows and cells/nuclei on the columns. **i.** Filter for low expressed genes (Methods: Expression and Proportion Zero filtering). Onward working with one cell type at time: **ii.** Compute Expression Rank of each cell/nucleus for each gene (example distribution in *b*), **iii.** Calculate mean gene expression across all cells/nuclei for one cell type and then its Rank Expression. **iv.** Per gene, find the difference of the Rank Expression against the mean Rank Expression for each cell/nucleus in a given cell type. **v.** Calculate the mean of the absolute Expression Rank differences for each gene. **vi.** Rank the mean absolute Expression Rank differences. **vii.** Repeat steps **ii-vi** for each cell type. **viii.** Per gene, compute the sum of the previous ranks across all cell types, and then rank these sums across genes such that the highest rank is given to the gene with the smallest sum. This is the final Rank Invariance value.

Using sc/snRNA-seq data, we define a candidate TREG as a gene that (1) has non-zero expression in most cells/nuclei across groups of interest, such as tissue-specific cell types, and (2) is expressed at a constant level with respect to other genes across different cell types of a given tissue. To be compatible with RNAscope, candidate TREGs also meet the following criteria: (3) expressed in the top 50% of genes for easy detection, (4) have a dynamic range of puncta to provide a continuous metric, (5) expressed at a level that individual puncta can be accurately counted.

While TREGs theoretically share some similarities with classical housekeeping (HK) genes, such as being expressed in every cell, they have other distinct properties. By definition, TREGs are tissue specific and are associated with total RNA expression. In other words, TREGs are identified in one reference dataset specific to an experimental condition; therefore TREGs are not necessarily generalizable to other experimental conditions. Furthermore, they are not defined by the function of the protein they encode. In contrast, classic HK genes are associated with cell maintenance, tissue agnostic, and expressed at a constant level regardless of cell type and condition [17].

While TREG is a general method, our research focus is motivated by understanding the transcriptional landscape in the human brain and identifying changes associated with psychiatric disorders [18]. We are interested in identifying a TREG that could be used in multiple cortical and subcortical brain regions linked to psychiatric disorders [18]. We focused on broad cell type categories that are diverse across size and expression levels, and are frequently present in these brain regions [18, 19]. With this in mind, we demonstrated the use of TREGs by applying our approach to snRNA-seq data from five brain regions, with focused RNAscope analyses in the dorsolateral prefrontal cortex (DLPFC). We compared candidate TREGs against classic HK genes and identified *AKT3* as the best performing TREG in the DLPFC. To identify candidate TREGs in other tissues, we provide open-source software available as an R/Bioconductor package at https://bioconductor.org/packages/TREG.

## Results

### Overview of method to identify TREGs

Our approach to identify Total RNA Expression Genes (TREGs) was inspired by rank-invariance methods originally developed for microarrays that were used to identify stably expressed genes within normalization methods applied to unbalanced transcriptome data (or containing different amounts of RNA) [7–9]. Briefly, after applying a filter to remove lowly expressed genes in a given sc/snRNA-seq reference dataset, our approach compares the ranks of expression across cells/nuclei (rather than comparing the gene expression values themselves across cells of different sizes) and identifies genes that are consistently ranked (or high ‘Rank Invariance’) (**Figure 1b**). In our algorithm, to identify a TREG, we compared the stability of each gene’s Expression Rank within and across cell types to identify high Rank Invariant genes (**Figure 1c,** Methods: Rank Invariance calculation). Genes consistently expressed in all cells/nuclei across all cell types were identified by high Rank Invariance values, and were considered TREG candidates. We implemented our data-driven method in an open-source R/Bioconductor package (https://bioconductor.org/packages/TREG) [20] to identify candidate TREGs in any sc/snRNA-seq dataset. The package includes functionality for both gene filtering and Rank Invariance methods.

### Datasets and TREG Experiment Overview

We applied our method to identify TREGs in a publicly available snRNA-seq dataset from the human postmortem brain. Specifically, the dataset included 70,527 nuclei from eight donors across five brain regions [18]. We identified candidate TREGs among 10 broad cell types across these brain regions: amygdala (AMY), dorsolateral prefrontal cortex (DLPFC), hippocampus (HPC), nucleus accumbens (NAc), and subgenual anterior cingulate cortex (sACC) (Methods: snRNA-seq reference data, **Supplementary Table 1**). Gene expression from top candidate TREGs was measured with smFISH using RNAscope technology and compared to a classic housekeeping gene, *POLR2A [21]*, to evaluate TREG predictiveness of total RNA expression.

### Filtering genes from the snRNA-seq data in the postmortem human brain

To maximize detection compatibility with the RNAscope assay, the expression data was filtered for highly expressed genes, specifically the top 50% of the 23,038 genes in the snRNA-seq dataset, retaining 11,519 genes. Genes were also filtered to remove those with a high maximum Proportion Zero (ranges between 0 and 1) expression across all cell type and brain region combinations (Methods: Expression and Proportion Zero filtering). The Proportion Zero filtering process avoids rank ties in the downstream steps due to the high number of genes with no expression. A high Proportion Zero also suggested that a gene may not be observable in most nuclei in that population using RNAscope. Frequently, nuclei from a specific cell type and brain region combination had a high frequency of genes whose Proportion Zero exceeded 0.75 for common cell types including astrocytes, microglia, oligodendrocytes, oligodendrocyte precursor cells, excitatory and inhibitory neurons (**Figure 2a)** and for more rare cell types including endothelial, macrophages, mural, and T-cells (**Supplementary Figure 1**). After filtering the genes for a maximum Proportion Zero of less than 0.75 across all cell types and region combinations, 877 genes remained (3.8% out of the initial 23,038 genes). The classic housekeeping gene *POLR2A* showed a high Proportion Zero in many cell types across brain regions and did not pass this filtering step unlike *AKT3, ARID1B, MALAT1* (**Figure 2b, Supplementary Figure 2a**).

**Figure 2:**
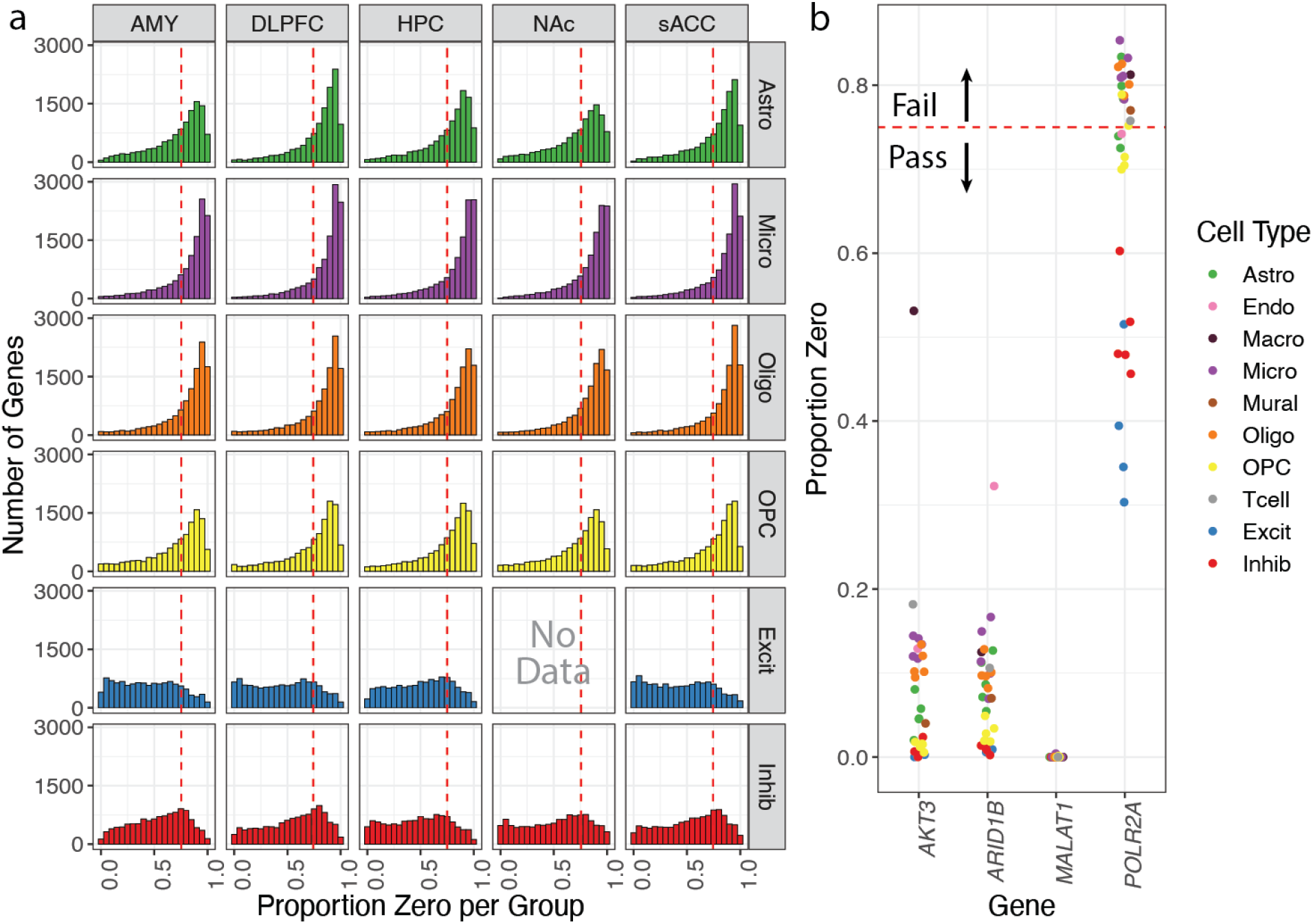
Overview of Proportion Zero filtering process. **a.** Histogram frequency of Proportion Zeros for each nuclei population for a given cell type and brain region combination. These combinations are arranged with cell types along the rows [astrocytes (Astro), microglia (Micro), oligodendrocytes (Oligo), oligodendrocyte precursor cells (OPC), excitatory (Excit) and inhibitory neurons (Inhib)] and by brain region along the columns [amygdala (AMY), dorsolateral prefrontal cortex (DLPFC), hippocampus (HPC), nucleus accumbens (NAc), and subgenual anterior cingulate cortex (sACC)]. Consistent with the inhibitory neuron-rich cell type composition of the NAc, there were no excitatory neurons found in this region and therefore no data to report. The red dashed line represents the 0.75 cutoff for filtering. **b.** Proportion Zero filtering process detailed for *AKT3, ARID1B, MALAT1* compared to the classic HK gene *POLR2A*. If any cell type and brain region combination (individual colored points), has a Proportion Zero > 0.75, then the gene fails the filtering step. Unlike *AKT3, ARID1B*, and *MALAT1, POLR2A* fails Proportion Zero filtering.

### Identification of TREG candidates in the postmortem human brain

After applying the filtering steps, the Rank Invariance workflow (**Figure 1c**) was applied to the five brain regions in the postmortem human brain to identify candidate TREGs (Methods: Rank Invariance calculation). From the top ten Rank Invariance values, we selected three candidate TREGs (*AKT3*, *ARID1B*, and *MALAT1*) for further evaluation based on the commercial availability of RNAscope probes. *MALAT1* was the top Rank Invariance gene, and also the gene with the highest mean expression. The Expression Rank of these TREGs has a small variance across 70k nuclei (**Figure 3a)**, as well as within different cell types (**Figure 3b**). This is in contrast to the HK gene *POLR2A*, which shows a more variable Expression Rank distribution (**Figure 3b**). We note that this same relationship holds if we compare the distribution of log-transformed gene expression across cell types, which is more variable than using the Expression Rank distribution (**Supplementary Figure 3**).

**Figure 3:**
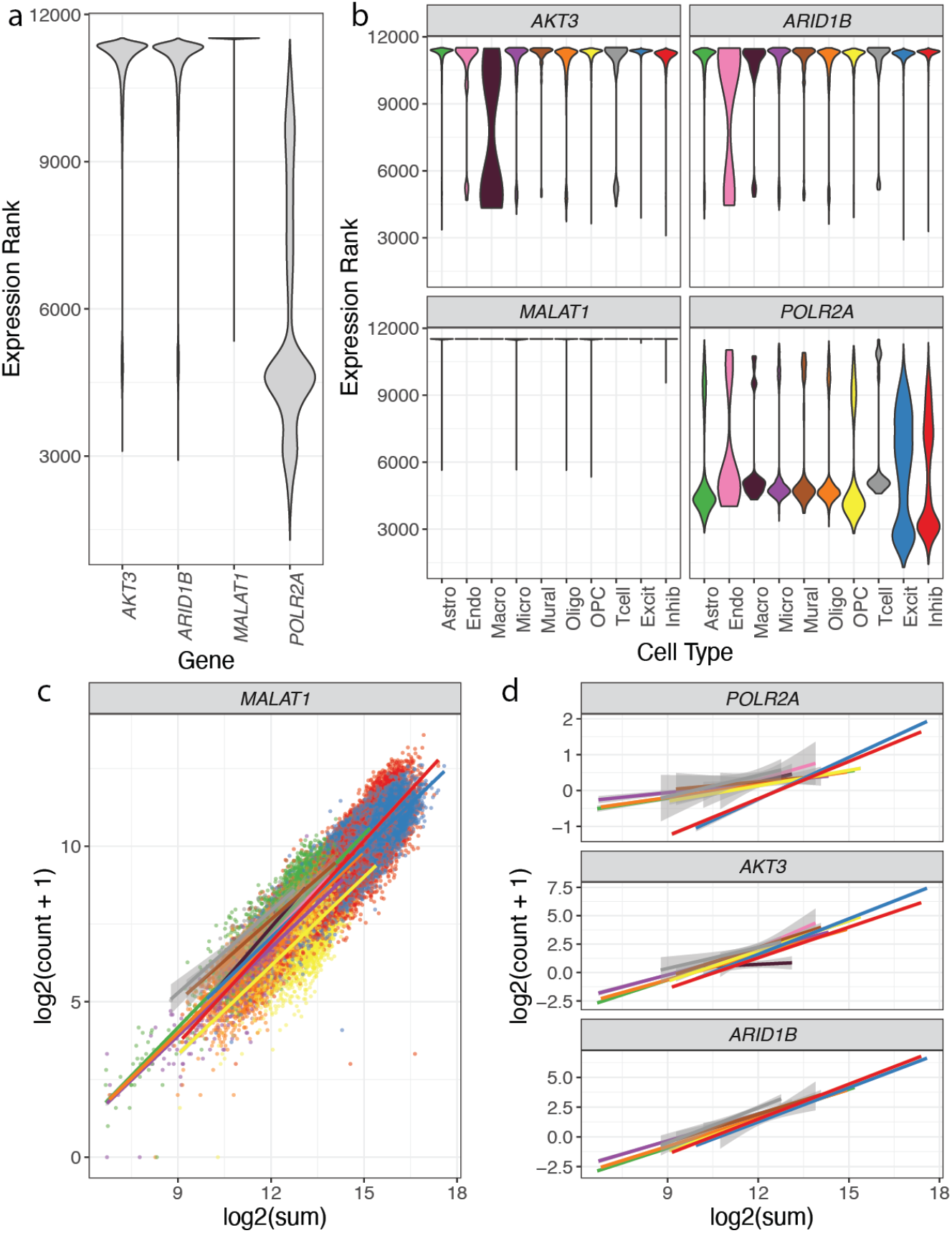
Distribution of ranks and relationship between total nuclear expression and expression of candidate TREGs. **a.** Distribution of the Expression Rank (*y*-axis) over all nuclei for genes *AKT3, ARID1B, MALAT1* (three candidate TREGs) and *POLR2A* (a known HK gene). The candidate TREGs show higher Rank Invariance compared to *POLR2A* (related to **Figure 1b**). **b.** The distribution of the Expression Ranks (*y*-axis) over all cell types (*x*-axis) for the three candidate TREGs shows less Expression Rank variability across most cell types compared to *POLR2A*. **c.** Scatter plot of the total RNA expression (estimated by the nuclei log2 sum of all counts) against the nuclei gene expression (log2 of the count plus one) for *MALAT1*, overlaid with the linear fit for each cell type and colored by cell type. **d.** Linear fits of total nuclear RNA expression against gene expression in the nuclei, similar to *c*. for *POLR2A, AKT3*, and *AR1D1B*. The expression of candidate TREGs show consistent positive linear relationships with total RNA expression in each nucleus across all cell types, unlike *POLR2A* where the neurons have a different pattern than other cell types.

By definition, the expression of a TREG should be predictive of the total expression of RNA in a cell(or nuclear expression when limited to snRNA-seq data). We compared the relationship between the gene expression of a TREG and the total expression of RNA in a nucleus (estimated by the log2 sum of all counts) and quantified the strength of the association by fitting a linear model for all nuclei within each cell type. We found a strong linear relationship for *MALAT1* (**Figure 3c**), *AKT3* and *ARID1B* (**Figure 3**, **Supplementary Figure 4**, **Supplementary Table 2**, Methods: Total RNA linear regression). Among the genes passing the Proportion Zero filter, the strength of their association with total RNA expression generally increased as their Rank Invariance increased (**Supplementary Figure 2b**). Furthermore, gene ontology enrichment analysis with the top 10% candidate TREGs showed that these genes are enriched for key biological processes such as RNA splicing and cellular components related to transcription (**Supplementary Figure 5**, Methods: Gene ontology enrichment analysis).

We ran the filtering process and TREG candidate identification independently for each of the five brain regions and identified the top 50 Rank Invariance genes (**Supplementary Table 2**). We identified 13 TREGs common across all five brain regions, therefore for the main analysis we used the combined dataset (**Supplementary Figure 6**). The top 13 TREGs across brain regions included, *AKT3, ARID1B*, and *MALAT1*.

### Validation of TREGs with smFISH using RNAscope technology

Next, we chose to further evaluate TREGs in DLPFC tissue given its implication in several psychiatric disorders. We used multiplex fluorescent smFISH with RNAscope technology to label candidate TREGs *AKT3, ARID1B*, and *MALAT1* as well as HK gene *POLR2A* in different cell types in DLPFC tissue sections from an independent brain donor (n=9 tissue sections with 3 tissue sections per gene, **Figure 4a**, Methods: Postmortem human tissue). We performed RNAscope with three probe combinations (**Supplementary Table 3-5**, Methods: RNAscope multiplex single molecule fluorescent in situ hybridization [smFISH]). Each combination probed a TREG with a panel of cell type marker genes including *SLC17A7, GAD1*, and *MBP* (labeling excitatory neurons, inhibitory neurons, and oligodendrocytes, respectively). *POLR2A* and *MALAT1* were hybridized in the same experiment, and due to limitations in multiplexing, *GAD1* was omitted. Following high magnification imaging, *AKT3, ARID1B*, and *POLR2A* transcripts were detected as discrete puncta (white dots) within individual nuclei (**Figure 5a-c**, Methods: Image acquisition). However, due to high expression, individual puncta could not be observed for *MALAT1* and fluorescent signals were too saturated for quantification (**Figure 5d**).

**Figure 4:**
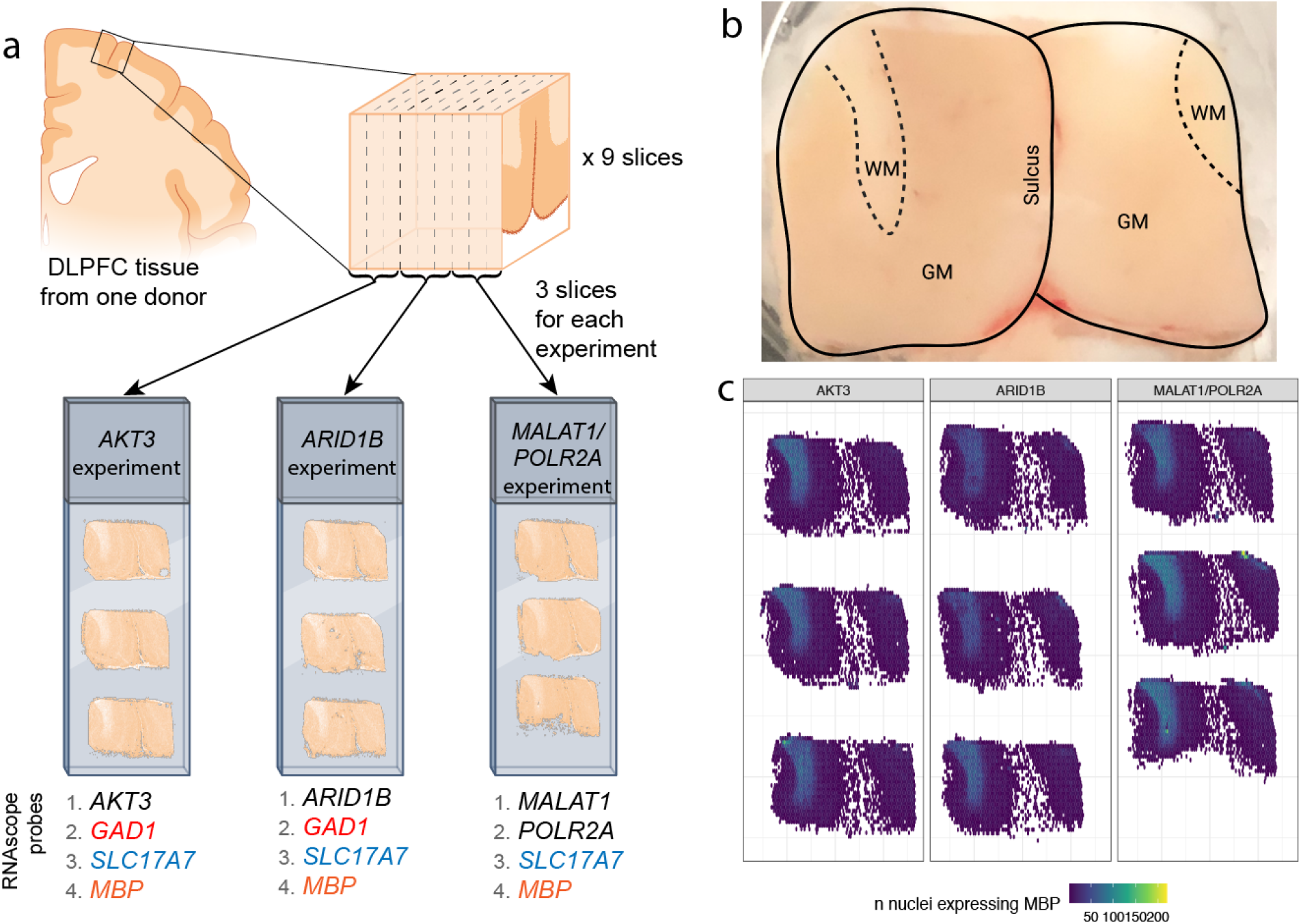
Overview of smFISH RNAscope experiment and DLPFC anatomy. **a.** Illustration of RNAscope experimental design where a single DLPFC tissue block was used to generate 9 spatially-adjacent slices. These 9 slices were hybridized with 3 RNAscope probe combinations noted as the *AKT3, ARID1B*, and *MALAT1/POLR2A* experiments (related to **Supplementary Table 3-5**). Candidate TREGs and *POLR2A* are shown in black, while *GAD1, SLC17A7*, and *MBP* are cell type marker genes for inhibitory neurons (red), excitatory neurons (blue), and oligodendrocytes (orange), respectively. **b.** Annotated image of DLPFC tissue, noting the location of gray matter (GM), white matter (WM), and sulcus. **c.** Spatial distribution of cells expressing *MBP* for each sample. *MBP* is an oligodendrocyte cell type marker gene enriched in white matter.

**Figure 5:**
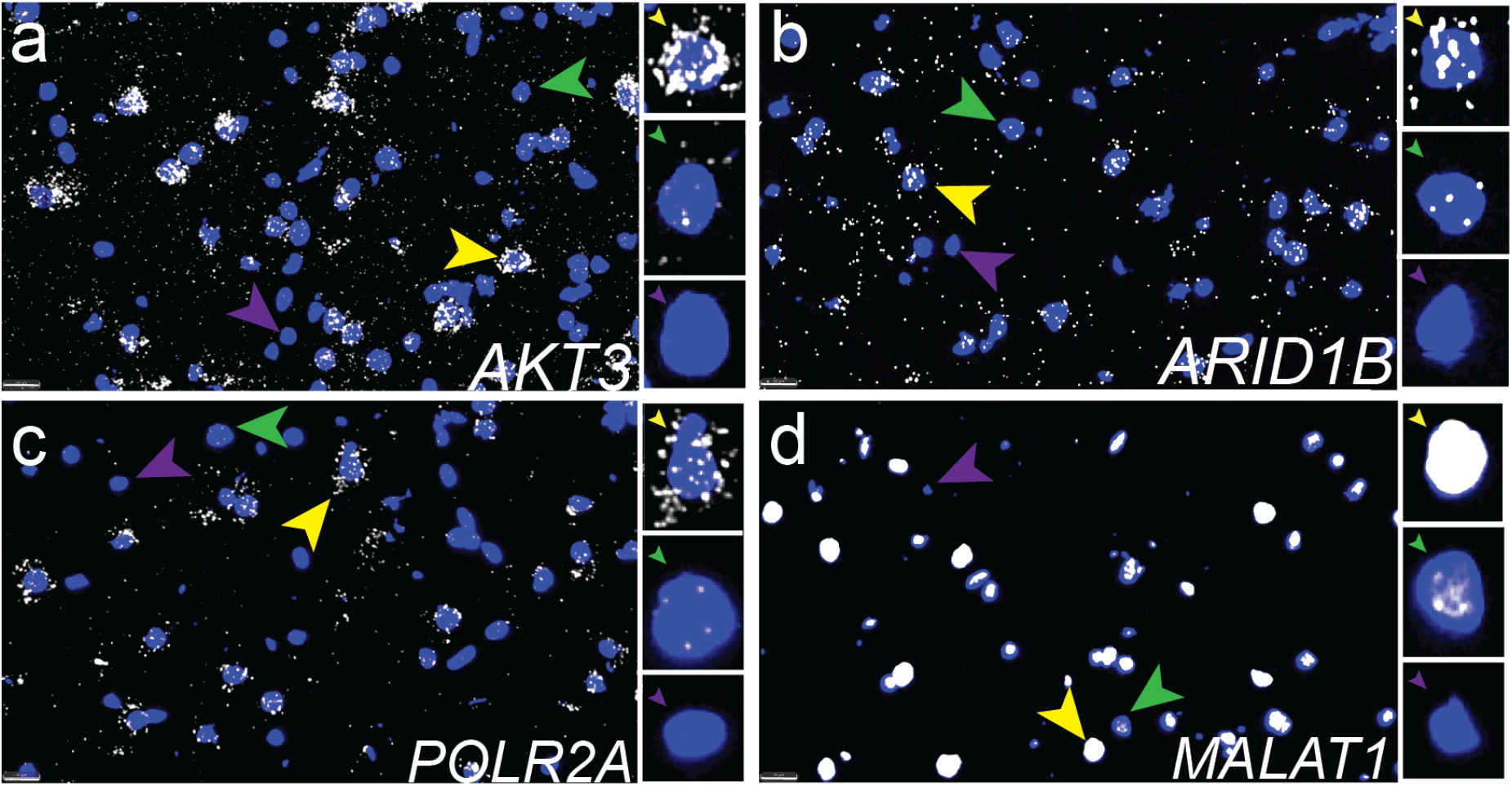
Expression of TREGs in individual nuclei using smFISH with RNAscope. Representative high magnification images showing expression of candidate TREGs **a.** *AKT3*, **b.** *ARID1B*, **c.** HK gene *POLR2A*, and **d.** *MALAT1* and in human DLPFC. Insets show individual nuclei with high expression (yellow arrow), low expression (green arrow), or in rare cases (<=14% for candidate TREGs and 22% for *POLR2A*, **Table 1**), no expression (purple arrow). Each puncta represents a single transcript, as illustrated in **Figure 1a**. *MALAT1* shows =extremely high expression in the majority of nuclei such that individual puncta cannot be quantified (yellow arrow). Scale bar is 20 um.

For TREGs showing discrete puncta, image segmentation and transcript quantification was performed using HALO software (Methods: Image analysis with HALO). HALO identified 1,099,931 individual nuclei across the nine DLPFC tissue sections, with 80k-109k nuclei segmented per tissue section (**Supplementary Table 6**). After quality control for poorly segmented regions (Methods: Quality control and spatial quantification of HALO data, **Supplementary Figure 7**) the number of nuclei per section was reduced to 68k-106k (**Supplementary Table 6**). We show accurate segmentation of fluorescent signals in a representative DLPFC tissue section including neuron-enriched gray matter and glial-enriched white matter (**Figure 4b-c**, **Figure 6**). As expected, quantification of nuclear area based on DAPI signal confirmed that neurons located in the gray matter have a larger nuclear area than glia located in the white matter (**Figure 6a, Supplementary Figure 7**). Quantification of *AKT3* puncta within nuclei confirmed higher expression of AKT3 in neuron-enriched gray matter, which is consistent with higher transcriptional activity in neurons compared to glia (**Figure 6b**). *ARID1B* and *POLR2A* also showed elevated expression in neurons located in the gray matter across the 3 different tissue sections (**Supplementary Figure 8**). Segmentation of *SLC17A7* (excitatory neurons), *GAD1* (inhibitory neurons), and *MBP* (oligodendrocytes) fluorescent signals revealed the expected spatial distribution of these cell types within the gray and white matter (**Figure 6c**).

**Figure 6:**
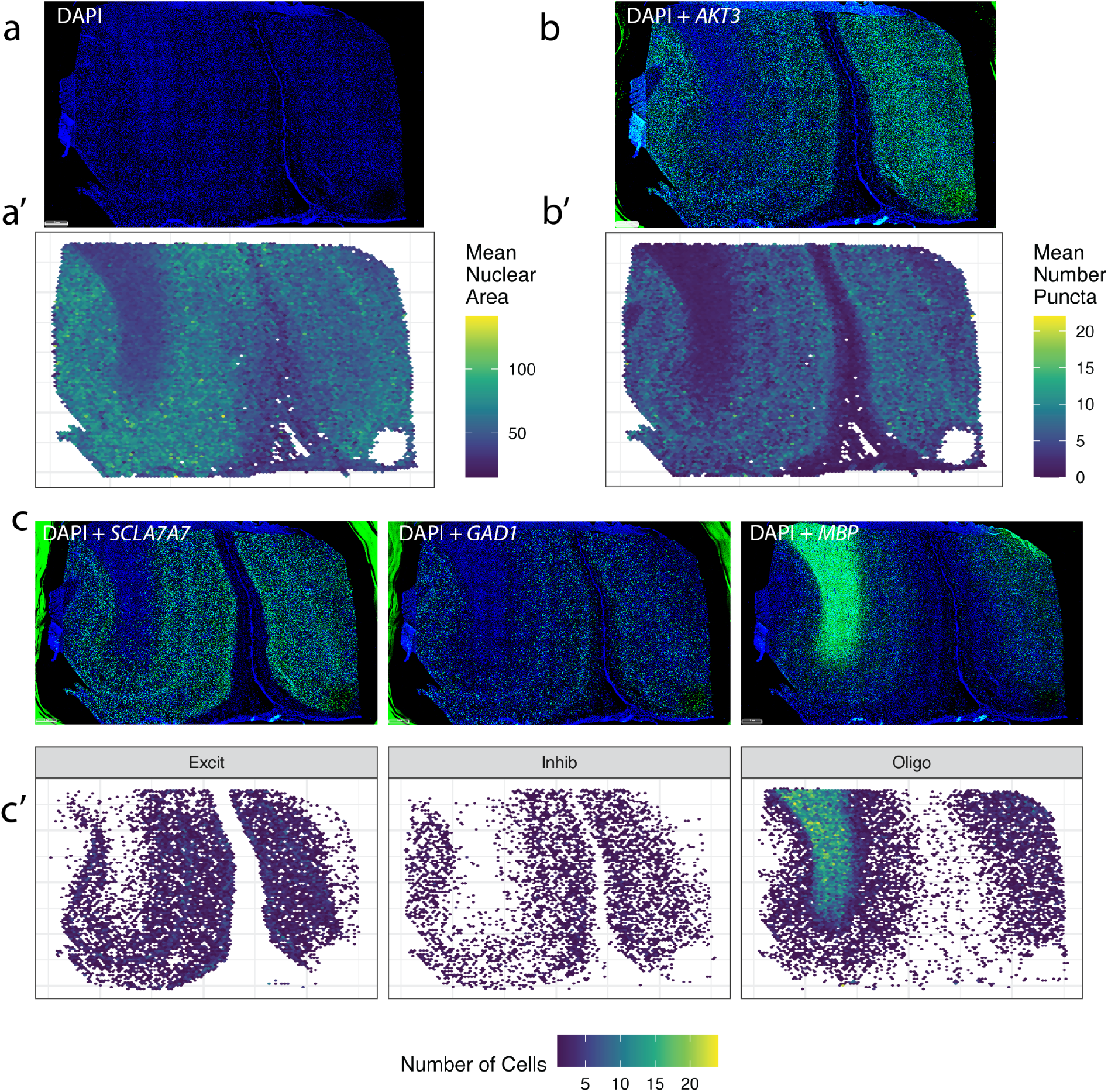
Quantification of candidate TREG *AKT3* in differently sized cell types in human DLPFC. Representative tissue section showing: **a.** Raw fluorescence for nuclear DAPI signals and **a’.** corresponding mean nuclear area size. Nuclear area based on DAPI signal shows larger excitatory neuron nuclei in gray matter from smaller glial nuclei in the white matter, related to **Figure 4b**. **b.** Raw fluorescence for DAPI and *AKT3* and **b’** corresponding quantification of mean number of *AKT3* puncta per nucleus. **c.** Raw fluorescence for *DAPI* and one of the following (*SLC17A7, GAD1*, and *MBP*) compared to **c’** the quantified distribution of the number of *SLC17A7+* excitatory neurons (Excit), *GAD1+* inhibitory neurons (Inhib), and *MBP+* oligodendrocytes (Oligo), respectively. Scale bar in a for a-c is 1 mm.

Image segmentation and transcript quantification revealed that candidate TREGs were consistently expressed in the majority of nuclei across cell types. Specifically, TREG expression was recorded in over 86% of nuclei (**Table 1**). The HK gene *POLR2A* had puncta that could be quantified in 78% of nuclei with RNAscope, which was unexpected given that only 30% of nuclei had non-zero expression values in snRNA-seq. Additionally, *AKT3 and ARID1B* provided a larger dynamic measurement range given that the mean puncta per nucleus is 4.09 and 2.08 respectively, compared to 2.75 for *POLR2A* (**Table 1**).

**Table 1:** Proportion of nuclei that displayed any TREG candidate or *POLR2A* puncta. Proportion of nuclei with a non-zero count in the DLPFC snRNA-seq data compared against the mean proportion of non-zero puncta in the nucleus and mean number of puncta observed in the RNAscope data for the candidate TREGs and *POLR2A*. Beta values are the slope of the linear fit of the number of puncta over ordered cell types and the 95% confidence interval. The standardized beta is the slope of the linear fit of the number of puncta divided by the standard deviation of the number of puncta for each gene. Standardized betas enable the comparison between snRNA-seq and RNAscope data. The standardized beta in snRNA-seq is −1.33 (−1.35,−1.31). With RNAscope, *AKT3* is the TREG that most similarly follows the trend across all genes in snRNA-seq (see also **Figure 7**). Due to the inability to resolve individual punca for *MALAT1*, the observed trend (**Supplementary Figure 9**) is unreliable.

| Gene | Prop. non-zero in DLPFC snRNA | Mean prop. non-zero puncta in the nucleus | Mean *n* puncta | $\beta$ (95% CI) | Standard deviation | Standardized $\beta$ (95% CI) |
| --- | --- | --- | --- | --- | --- | --- |
| *AKT3* | 0.92 | 0.88 | 4.09 | -5.52<br>(-5.55,-5.49) | 5.18 | -1.07<br>(-1.07,-1.06) |
| *ARID1B* | 0.94 | 0.86 | 3.08 | -2.63 (-2.65,-2.6) | 3.42 | -0.77<br>(-0.77,-0.76) |
| *MALAT1* | 1.00 | 0.98 | 2.07 | -1.22<br>(-1.24,-1.21) | 1.53 | -0.8<br>(-0.81,-0.79) |
| *POLR2A* | 0.30 | 0.78 | 2.75 | -3.49<br>(-3.51,-3.47) | 3.34 | -1.05<br>(-1.05,-1.04) |
| *All genes in snRNA-seq* | NA | NA | NA | -21844.07<br>(-22172.45,-21515.68) | 15560.76 | -1.33<br>(-1.35,-1.31) |

Next, we evaluated how the number of measured puncta (for each TREG) in a nucleus related to total nuclear RNA expression measured by snRNA-seq for excitatory neurons, inhibitory neurons, and oligodendrocytes in the DLPFC. DLPFC snRNA-seq data showed that excitatory neurons have the highest total nuclear RNA expression (estimated with the sum of total UMI counts per nucleus), followed by inhibitory neurons, and then oligodendrocytes (**Figure 7a**). We quantified this pattern using the standardized regression coefficient for total nuclear RNA expression vs. cell types, which is −1.33 (95% CI = −1.35,−1.31, **Table 1**, Methods: Linear regression of puncta across cell types). For candidate TREGS *AKT3* and *ARID1B*, we measured the number of puncta per nucleus across cell types and found that *AKT3* tracks the closest with the pattern of total RNA expression measured by snRNA-seq and has a more symmetric distribution than *POLR2A*, particularly for oligodendrocytes (**Figure 7b**). Excitatory neurons contain the most puncta, followed by inhibitory neurons, and then oligodendrocytes.

**Figure 7:**
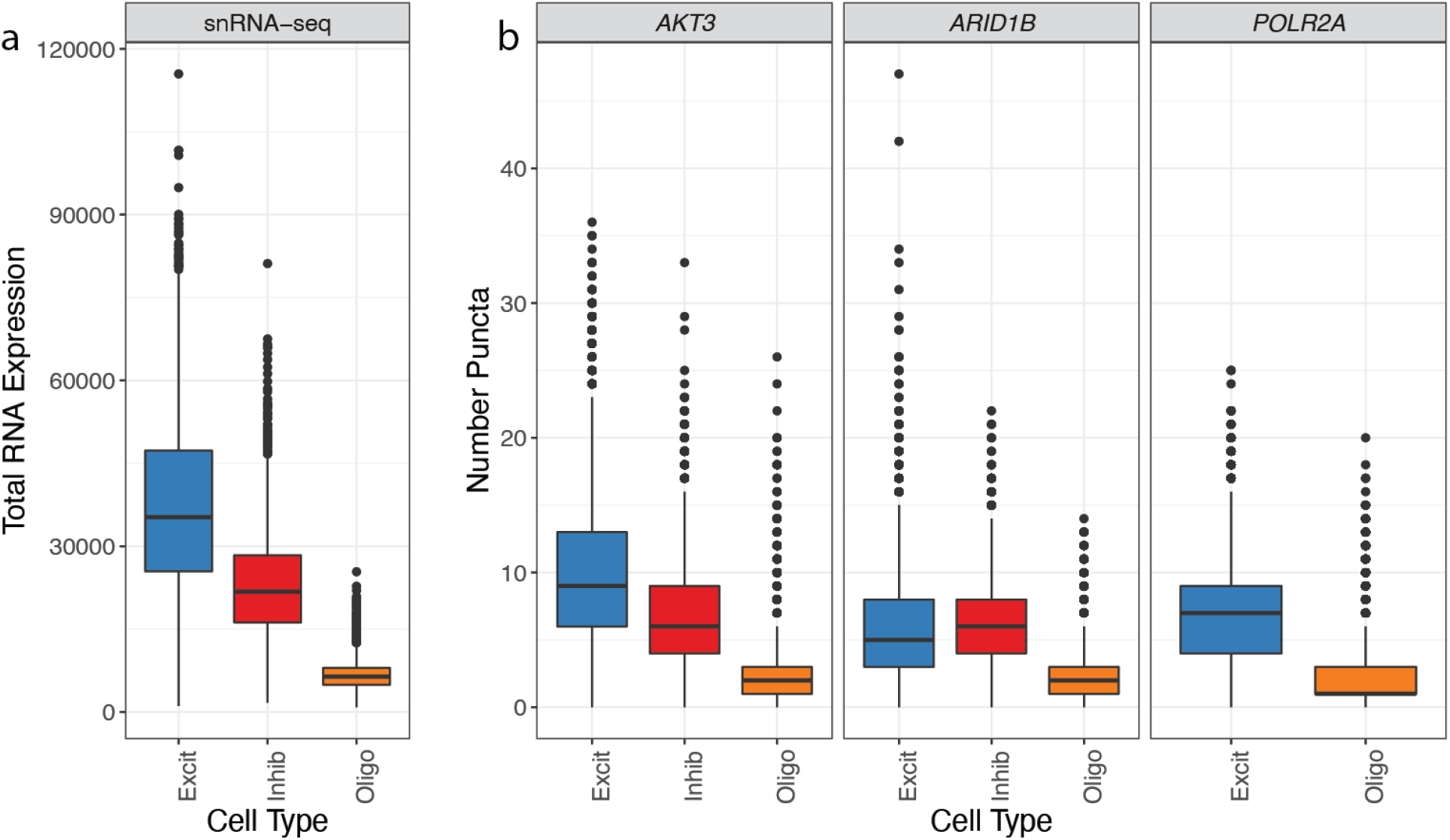
Boxplots of total RNA nuclear expression in the nucleus across cell types. **a.** Distribution of total nuclear RNA expression (estimated with the sum of total UMIs per nucleus) in DLPFC snRNA-seq data across excitatory neurons (Excit), inhibitory neurons (Inhib), and oligodendrocytes (Oligo). **b.** Distribution of the number of puncta quantified by RNAscope for each observed gene across the same cell types as in *a*. The number of puncta by RNAscope estimates the total RNA expression by snRNA-seq (**Figure 1a)**. *POLR2A* was only evaluated in excitatory neurons and oligodendrocytes as it was multiplexed with *MALAT1* and and *GAD1* was omitted.

*ARID1B* breaks from the expression pattern shown in the reference snRNA-seq data, given that inhibitory neurons are higher than excitatory neurons, although neurons still show more expression than the oligodendrocytes. *POLR2A* was only measured in two cell types, excitatory neurons and oligodendrocytes, but also follows this pattern with higher expression in neurons compared to glia. The standardized regression coefficient for number of puncta vs. cell types for *AKT3* is −1.07 (95% CI = −1.07,−1.06) and is the closest to the snRNA-seq coefficient (−1.33, 95% CI = −1.35,−1.31) of the experimentally validated genes (**Table 1**). This pattern is also consistent when considering other cell types (**Supplementary Figure 9**). By measuring total nuclear RNA abundance and nuclear area across cell types in the same experiment, we demonstrate that the relationship between RNA content and nuclear area changes across cell types (**Supplementary Figure 10**).

## Discussion

In this work, we showed that our new data-driven Rank Invariance method successfully determines candidate total RNA expression genes (TREGs) from snRNA-seq data that can be used in combination with smFISH to accurately estimate relative RNA abundance in distinct cell types of varying sizes and transcriptional activity. We investigated the properties of TREGs by evaluating the consistency of the Expression Ranks and relationship with total RNA expression in snRNA-seq data. We further validated TREG candidates by quantifying puncta with smFISH using RNAscope, which found that *ATK3* best reconstructed the pattern between cell type and total RNA expression observed in human DLPFC snRNA-seq data. While the Rank Invariance method was successful, it cannot guarantee that candidate TREGs will be experimentally compatible for downstream needs. For example, *MALAT1* was the top candidate TREG in our study, but *MALAT1* was incompatible with resolving individual puncta by RNAscope because of its extremely high expression. 10x Genomics advises users that independent of protocol *MALAT1* is frequently observed in poly-A captured RNA-seq data [22], which is consistent with comparisons between polyA selection versus rRNA depletion protocols [23] and could be due its structure [24]. Furthermore, *MALAT1* has been used as a proxy for nuclear expression linked to damaged cells in scRNA-seq data [25]. We thus recommend that TREGs be evaluated in the assay or experimental setting of choice before implementing experiments using rare and valuable samples.

To identify relevant TREGs for a particular study, it is important to use sc/snRNA-seq data that is compatible with the experimental design. That is, sc/snRNA-seq data should contain the tissue and cell types combinations of interest, as well as match the experimental conditions for which the TREG will be used, such as age, sex, diagnosis, and experimental exposure. Otherwise candidate TREGs might be less reliable for quantifying total RNA as they could be specific to an organism, tissue, cell type, or experimental condition. As more snRNA-seq datasets come online across tissues and experimental organisms, our Rank Invariance methodology can be used to identify TREGs in different experimental settings. Furthermore, the mean absolute differences (**Figure 1c** arrow **v.**) will be more stable when larger numbers of cells/nuclei are present per cell type. Thus, the Rank Invariance process might be less reliable for datasets with rare cell types, for which it might be best to perform a sensitivity analysis without the rare cell types to compare resulting TREGs and ultimately identify reliable TREGs.

While TREGs share some similarities with housekeeping (HK) genes, they are by definition different. A HK gene typically has a defined central molecular function and is expected to be expressed at a constant level across multiple tissues [17]; for example the GTEx Portal [26] shows less variable expression across tissues for *POLR2A* than *AKT3* (https://gtexportal.org/home/gene/POLR2A vs https://gtexportal.org/home/gene/AKT3). In contrast, the RNA level of a TREG should be quantifiable in most cells among all cell types in a particular experimental setting, and most importantly, it should be highly predictive of the total RNA expression of those cells or nuclei. In our snRNA-seq data, *POLR2A* and other HK genes had high Proportion Zero and were not as strongly predictive of total nuclear RNA expression as other candidate TREGs. Interestingly, by RNAscope, *POLR2A* could be measured in a larger percent of nuclei compared to snRNA-seq (78% vs 30%, **Table 1**). We note that TREGs were detected in the majority of nuclei by RNAscope as expected, but we did observe some nuclei lacking expression. This could be due to only a fraction of the nucleus being present in the 10um tissue section plane or technical limitations in multiplex fluorescent slide scanning with spectral imaging, including low resolution and image acquisition in only the x and y dimensions, with no z axis component [27]. However, *AKT3* was present in 88% of nuclei by RNAscope and had a higher mean number of puncta compared to *POLR2A* (4.09 vs 2.75). *AKT3* best recapitulated the observed trend in snRNA-seq data (**Figure 7b**). Thus while *POLR2A* performed better than expected on RNAscope, *ATK3* still outperformed *POLR2A* across different metrics.

The protein encoded by *AKT3* is a member of the AKT/protein kinase B family of serine/threonine protein kinases. AKT plays a key role in the phosphatidylinositol 3-kinase (PI3K)-Akt-mTOR signaling cascade, which regulates numerous biological processes such as cell proliferation, growth, apoptosis, and metabolism [28]. *AKT3* is one isoform of AKT that is predominantly expressed in the human and mouse brain and plays a significant role in brain development [29]. Dysfunction of AKT3 is implicated in a variety of neurodevelopmental and neurodegenerative brain disorders and tumors, such as glioma [29, 30]. The *AKT3* gene has also been associated with risk for neuropsychiatric disorders [31]. AKT3 is an important enzyme whose function needs to be carefully regulated at the protein level. Thus, *AKT3* gene expression is likely highly regulated across cell types, which is consistent with its dynamic expression in neurons and glia (**Supplementary Figure 3**). While candidate TREGs, such as *AKT3* and *ARID1B*, are clinically-relevant [27, 29–34], the snRNA-seq data used in this study is from neurotypical control subjects. As more snRNA-seq datasets are generated from subjects with neuropsychiatric and neurological disorders, it will be important to assess candidate TREGs in the context of brain disorders. More generally, other candidate TREGs identified in our neurotypical control tissue are functionally implicated in transcription (**Supplementary Figure 5**), which is consistent with the notion that a TREG should be predictive of total RNA expression.

In contrast to rank-invariant methods [7–9], the method we developed here is not a normalization method, but a method for gene selection that is specific to the desired properties of a TREG, namely correlation with total RNA expression. Our implementation is thoroughly tested with 100% code coverage [35] and available via Bioconductor at https://bioconductor.org/packages/TREG [20]. While our list of candidate TREGs could be valid for other DLPFC studies (**Supplementary Table 2**), we recommend that you use our R package with your own sc/snRNA-seq data. TREGs could be useful for other research purposes and other contexts than the ones envisioned here.

We used smFISH with RNAscope technology to validate candidate TREGs across three major cell types in the human DLPFC: oligodendrocytes, excitatory neurons, and inhibitory neurons. We selected only 3 cell types due limitations in multiplexing with RNAscope (the V2 assay supports a maximum of 4 targets). In the future, we plan to expand our experimental design to include other major cell types captured in snRNA-seq data, such as astrocytes, microglia, and oligodendrocyte progenitor cells. Another limitation of our study is that we focused only on TREG expression in the nucleus, but many TREGs are also expressed in the cytoplasm. Currently, the HALO FISH-IF module will only quantify puncta within a nucleus or a dilated area around the nucleus, which is a limitation when working with cell types that are not oval in shape, such as neurons and glia. While our analysis was focused only in the nucleus, previous work suggests that gene-level expression between the nucleus and cytoplasmic compartments are comparable [16, 36]. In future studies, we aim to improve cell segmentation to be able to estimate cell size and RNA content instead of restricting analyses to the nucleus.

While sc/snRNA-seq and bulk RNA-seq data is commonly derived from pulverized tissue, our work suggests considering a different experimental design. In particular, if you are designing a paired sc/snRNA-seq and bulk RNA-seq study where you will use the snRNA-seq data as a reference for deconvolution of bulk RNA-seq, generating spatially-adjacent dissections in order to use them for RNAscope experiments could be useful to “future-proof” your datasets. With this experimental design, you could identify cell types in the sc/snRNA-seq data, then identify candidate TREGs based on those cell types, and use these candidate TREGs with smFISH to quantify size and total RNA content for the cell types of interest (**Supplementary Figure 10**). These cell metrics could improve the precision of deconvolution algorithms and generate a potential gold standard dataset to evaluate the performance of the deconvolution methods.

## Conclusion

Through the data-driven Rank Invariance process, we have identified several candidate genes as Total RNA Expression Genes (TREGs) in five postmortem human brain regions. RNAscope validation experiments revealed that *AKT3* is a strong proxy for total nuclear RNA expression in the DLPFC. Future work will use individual TREGs to estimate total RNA abundance in differently sized cell types of the DLPFC to bolster deconvolution approaches. As more snRNA-seq data comes online, this Rank Invariance methodology could facilitate identification of TREGs in other experimental settings with differences in donor demographics, brain regions, tissues, or species. Similar to highly variable genes or housekeeping genes, TREGs represent an important class of genes that could be used for a variety of assays and downstream analyses. Our method for identifying TREGs is accessible, integrated with the Bioconductor ecosystem [37], and available at https://bioconductor.org/packages/TREG [20].

## Methods

R [38] and HALO (version 3.3.2541.383, Indica Labs) was used for the data analysis, and plotting was done with the ggplot2 [39] and UpSetR [40] packages.

### snRNA-seq reference data

The single nucleus RNA-sequencing (snRNA-seq) reference data used for the Rank Invariance process is a publicly available dataset (10x Genomics, single cell 3’ gene expression) from postmortem human brain, which includes tissue from eight donors and five brain regions, including the amygdala (AMY), dorsolateral prefrontal cortex (DLPFC), hippocampus (HPC), nucleus accumbens (NAc), and subgenual anterior cingulate cortex (sACC) [18]. The original study classifies many region-specific subtypes of inhibitory and excitatory neurons, however for the purpose of this study, a lower resolution of broad cell types was used: astrocytes (Astro), endothelial cells (Endo), microglia, mural cells, oligodendrocytes (Oligo), oligodendrocyte precursor cells (OPC), T-cells, excitatory (Excit) and inhibitory neurons (Inhib). Specialized medium spiny neurons found exclusively in the NAc were classified as Inhib (**Supplementary Table 1**).

### Expression and Proportion Zero filtering

In R [38], by default, the `rank ()` function returns high ranks for high values, where equivalent values or ties are given an average value. To reduce the occurence of ties, we removed genes that would introduce many low-value ties. The data was first filtered to the top 50% of expressed genes. The nuclei were grouped by cell type and brain region, the Proportion Zero counts was calculated for each gene in each group, and is defined as 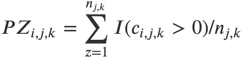 where 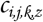 is the number of snRNA-seq counts for cell/nucleus *z* for gene *i*, cell type *j*, and brain region 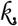 and 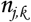 is the number of cells/nuclei for cell type *j* and brain region 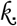 In our dataset, if the cell type was rare (less than or equal to 100 total nuclei, as was the case for Endo, Macro, Mural, and T-Cells) the nuclei from different regions were combined into one group, effectively ignoring the brain region from which the cell type was derived (**Supplementary Figure 1**). The distribution of Proportion Zeros for each group was visualized and used to select a cutoff value of 0.75, which included the peak of the Proportion Zero distributions (**Figure 2a**). Then for each gene, the maximum Proportion Zero across groups was required to be less than the cutoff (i.e. < 0.75) to pass the filtering step (**Figure 2b)**.

### Rank Invariance calculation

After Proportion Zero filtering, the remaining genes were evaluated for Rank Invariance jointly across all five brain regions, thus the nuclei were grouped only by cell type. The normalized expression (`logcounts`) of each gene was ranked for each nucleus, and the result was a matrix of Expression Rank values (the number of nuclei * number of genes). Within each cell type, the mean expression for each gene was ranked, and the result was a vector called mean Expression Rank (length is the number of genes). Then, the absolute difference between the Expression Rank of each nucleus and the mean Expression Rank was found. From here, the mean of the differences for each gene was calculated and then ranked. These steps were repeated for each cell type, and the result was a matrix of ranks, (number of cell types * number of genes). From here, the sum of the ranks for each gene were reverse ranked such that low values were given a high rank. This process resulted in the final value for each gene called the Rank Invariance value (**Figure 1b**). The genes with the highest Rank Invariance were considered as candidate TREGs. Classic housekeeping (HK) genes [41, 42] and brain data-driven HK genes [6] that fail these filtering steps are shown in **Supplementary Figure 2**.

### Total RNA linear regression

We tested for an association between the expression of each gene and the overall RNA expression of each nucleus using a linear regression model `log2 (counts + 1) ~ log2 (sum) + cellType` with limma package version 3.48 and “voom” [43, 44]. (**Figure 3c-d**). The *t*-statistics from this analysis are plotted in **Supplementary Figure 2**. The combination of the Rank Invariance values and the rank of the *t*-statistics from this linear model were used to help identify the best candidate TREGs.

### Gene ontology enrichment analysis

Of the 877 of genes evaluated for Rank Invariance, the top 10% were evaluated for gene ontology enrichment. Of the 23,038 genes in the snRNA-seq dataset, 18,296 have entrez ids and were used as the universe, of the 87 top RI genes 86 have entrez ids and were provided as the sole gene cluster. The enrichment analysis was performed with the `compareCluster ()` function from clusterProfiler package version 4.2 [45, 46]. Ontologies biological processes (BP), cellular components (CC), and molecular function (MF), were all tested.

### Postmortem human tissue

The human postmortem brain used in this study for RNAscope was obtained by autopsy from the Offices of the Chief Medical Examiner of the District of Columbia and of the Commonwealth of Virginia, Northern District, with informed consent from the legal next of kin (protocol 90-M-0142 approved by the NIMH/NIH Institutional Review Board). Details regarding curation, diagnosis, tissue handling, processing, and quality control measures have been described previously [47]. The study included a single neurotypical control adult donor (Br1531). A small piece of frozen DLPFC was dissected under visual guidance with a handheld dental drill on dry ice by a neuroanatomist. Gray and white matter tissue from the crown of the middle frontal gyrus was obtained from the coronal slab corresponding to the middle one-third of the DLPFC (along its rostral-caudal axis) immediately anterior to the genu of the corpus callosum. Microdissected DLPFC tissue was stored at −80°C until cryosectioning.

### RNAscope multiplex single molecule fluorescent in situ hybridization (smFISH)

DLPFC tissue was cryo-sectioned at 10μm on a Leica cryostat. Three tissue sections were collected per slide. Prior to use, the slides were stored at −80°C. Using RNAscope technology (RNAscope Multiplex Kit V2 and 4-plex Ancillary Kit: Cat # 323100, 323120, Advanced Cell Diagnostics, Newark California), probe hybridization and labeling was completed following the manufacturer’s instructions. Briefly, the protocol includes fixing the tissue sections in 10% neutral buffered formalin solution (Cat # HT501128-4L, Sigma-Aldrich, St. Louis, Missouri), dehydration in a series of ethanol solutions, pretreatment with hydrogen peroxide, and permeabilizing with proteases. Each slide was then incubated with one of the following probe combinations: *AKT3/GAD1/SLC17A7/MBP; ARID1B/GAD1/SLC17A7/MBP; POLR2A/MALAT1/SLC17A7/MBP* (**Supplementary Table 3**). These combinations were named according to the candidate TREGs or housekeeping (HK) gene included (i.e. *AKT3, or ARID1B*, or *POLR2A/MALAT1*) (Cat # 434211, 404031-C2, 415611-C3, 411051-C4, 551721, 310451, 400811-C2, Advanced Cell Diagnostics, Newark California). After washing briefly, slides were stored in 4X saline-sodium citrate (Cat # 351-003-101, Quality Biological, Gaithersburg Maryland) overnight at 4°C. Probes were then fluorescently labeled using opal dyes (Opal 520, Opal 570, Opal 620, and Opal 690, Perkin Elmer, Waltham, MA). Dyes were assigned to probes and diluted in concentration as described in **Supplementary Table 3** and **Supplementary Table 4**. Nuclei were labeled with DAPI (4′,6-diamidino-2-phenylindole) and coverslipped with fluoromount-G mounting media.

### Image acquisition

Slides were imaged at 20x magnification using a Vectra Polaris Automated Quantitative Pathology Imaging System (Akoya Biosciences), which performs multi-spectral imaging. For each probe combination a scanning protocol was created. Each protocol optimized the exposure time for a given opal dye in each probe combination as listed in **Supplementary Table 5**. Scanning generated a large `.qptiff` image file, which was then pre-processed in Phenochart (Akoya Biosciences). Briefly, the boundary of each slide (including the 3 tissue sections) was traced, and the individual `.tiff` tiles making up the scan area (1141-1489 tiles per slide) were extracted. These tiles were then subjected to linear unmixing in InForm (Akoya Biosciences). Unmixed `.tiff` tiles were then fused in HALO (version 3.3.2541.383, Indica Labs).

### Image analysis with HALO

Fused images from each scanned slide were annotated in HALO by drawing a boundary around each tissue section. The annotated areas across tissue sections ranged from 156716789.46μm^2^ to 162640367.42μm^2^ and annotations were consistent among the tissue sections on each slide. The FISH-IF module (version 2.1.5) was then used to segment cells and assign phenotype (i.e. cell type). Briefly, we first assigned dyes to either FISH or immunofluorescence (IF). While these experiments were exclusively FISH, the distinction between FISH and IF dyes allow for visualization and segmentation of diffuse staining vs. individual puncta. DAPI, *GAD1, SLC17A7*, and *MBP* were assigned IF values given their strong signals resembling diffuse labeling. The FISH dye assignment changed for each experiment (*AKT3*, *ARID1B, POLR2A/MALAT1*). Segmentation was optimized for each dye for each tissue section by adjusting several values with reference to manufacturer’s guidelines: HALO 3.3 FISH-IF Step-by-Step guide (Indica labs, Version 2.1.4 July 2021) and Quantitative RNAscope Image Analysis Guide (Indica labs). Size thresholds for nuclei, cytoplasm, cells, and FISH probe puncta were held constant across all tissue sections. Once the puncta counting was completed, the object data and settings were exported as `.csv` files (available via Globus at ‘jhpce#TREG_paper’) and `.txt` files (available on Github), respectively.

### Quality control and spatial quantification of HALO data

Visual inspection of the images revealed some technical artifacts related to cryosectioning and slide scanning (**Supplementary Figure 7**), including tissue tears/shredding, small bubbles, and out of focus fields. Out of focus fields caused nuclei to appear bigger and blurred together multiple puncta so they were not clearly resolved. Nuclei from these regions were excluded from the data analysis during these quality control steps.

Nuclei with only *MBP* expression were classified as oligodendrocytes (Oligo), with only *SLC17A7* expression as excitatory neurons (Excit), and with only *GAD1* expression as inhibitory neurons (Inhib). Nuclei with expression of multiple marker genes that could not definitively be assigned a cell type were classified as “Multi”. Nuclei with no markers were classified as “Other” and likely represent other non-neuronal cell types in the brain that were not labeled, including astrocytes and microglia. As the *MALAT1/POLAR2A* samples were not labeled with *GAD1* due to technical limitations in multiplexing (**Supplementary Table 3**), the number of Inhib nuclei could not be determined for these samples (**Supplementary Table 6**).

### Linear regression of puncta across cell types

TREG candidates were evaluated by the proportion of cells where any puncta were recorded in the HALO segmented nuclear area, as well as the mean number of puncta recorded (**Table 1**). The pattern of expression across cell types was compared to the sum of total counts of that cell type in the reference snRNA-seq data (**Figure 7**). To quantify this relationship we estimated the regression coefficient of total RNA over the three cell types that were sampled (Excit, Inhib, Oligo), for the snRNA-seq total RNA of a nucleus was estimated by the sum of UMIs. For the RNAscope data, total nuclear RNA is estimated by the number of segmented puncta. To compare these different data types, the standardized regression coefficient was calculated by dividing by the standard deviation of the total UMIs and number of puncta respectively.

## Abbreviations

AMY: amygdala
Astro: astrocytes
DLPFC: dorsolateral prefrontal cortex
Excit: excitatory neurons
Expression Rank: rank of the log normalized counts expression values for a given gene and nucleus, with high expression values translating into high rank values
HPC: hippocampus
HK: housekeeping
Inhib: inhibitory neurons
Micro: microglia
NAc: nucleus accumbens
Oligo: oligodendrocytes
OPC: oligodendrocyte progenitor cells
Proportion Zero: defined in Methods: Expression and Proportion Zero filtering
RI: Rank Invariance
sACC: subgenual anterior cingulate cortex
smFISH: single-molecule fluorescent in situ hybridization
TREG: total RNA expression gene

## Declarations

### Ethics approval and consent to participate

Not applicable.

### Consent for publication

Not applicable.

### Availability of data and materials

The snRNA-seq data [18] is available from https://github.com/LieberInstitute/10xPilot_snRNAseq-human#processed-data [48]. The code for the analysis is available from https://github.com/LieberInstitute/TREG_paper [49] and the software from https://github.com/LieberInstitute/TREG [20]. The raw data are publicly available from the Globus endpoint ‘jhpce#TREG_paper’ that is also listed at http://research.libd.org/globus/jhpce_TREG_paper/. The raw data provided through Globus include all the raw image files and the HALO .csv segmentation results.

### Competing interests

The authors declare that they have no competing interests.

### Funding

This project was supported by the Lieber Institute for Brain Development, National Institutes of Health grant R01 MH123183 (LAHM, KDM, SHK, SCP, SCH, KRM, LCT), and CZF2019-002443 (SCH) from the Chan Zuckerberg Initiative DAF, an advised fund of Silicon Valley Community Foundation. All funding bodies had no role in the design of the study and collection, analysis, and interpretation of data and in writing the manuscript.

### Authors’ contributions

- LAHM: Conceptualization, Formal Analysis, Methodology, Software, Visualization, Writing - original draft, Writing - review & editing
- KDM: Investigation, Validation, Visualization, Writing - original draft
- SHK: Formal Analysis, Investigation, Validation, Visualization, Writing - original draft
- SCP: Writing - review & editing
- SCH: Conceptualization, Funding Acquisition, Writing - review & editing
- KRM: Conceptualization, Funding Acquisition, Project Administration, Supervision, Writing - original draft, Writing - review & editing
- LCT: Conceptualization, Methodology, Software, Supervision, Writing - original draft, Writing - review & editing

## Acknowledgements

While an Investigator at Lieber Institute for Brain Development (LIBD), Andrew E. Jaffe helped frame this project by conceptualizing the utility of TREGs. We thank Keri Martinowich for consultation on RNAscope experimental design and comments on the manuscript. Matthew N. Tran from LIBD provided details and insight on the snRNA-seq data used in this project, and offered feedback on methodology. We would also like to acknowledge Thomas M. Hyde and the LIBD Neuropathology team, particularly James Tooke, for their help with the brain tissue dissection used in the RNAscope experiments. We acknowledge Elizabeth Engle and the Tumor Microenvironment Lab core facility for assistance with the Vectra Polaris slide scanner.

## Supplementary Material

### Supplementary Figures

**Supplementary Figure 1:**
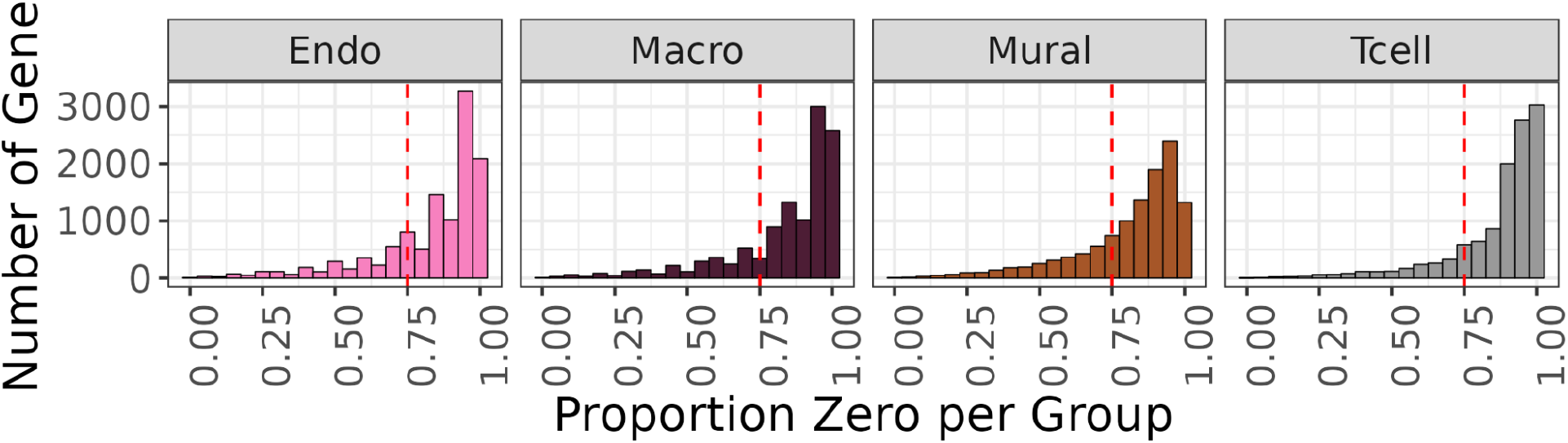
Distribution of Proportion Zero for rare cell types. Endothelial cell (Endo), Macrophage (Macro), Mural cell (Mural), and T-cell are cell types that have 100 (0.14%) or fewer nuclei in the snRNA-seq dataset used in this study (**Supplementary Table 1**). Given they are present in at most three of five brain regions, we did not include brain regions when grouping nuclei for the Proportion Zero filter calculation. Related to **Figure 2a**, Methods: *Expression and Proportion Zero Filtering*.

**Supplementary Figure 2:**
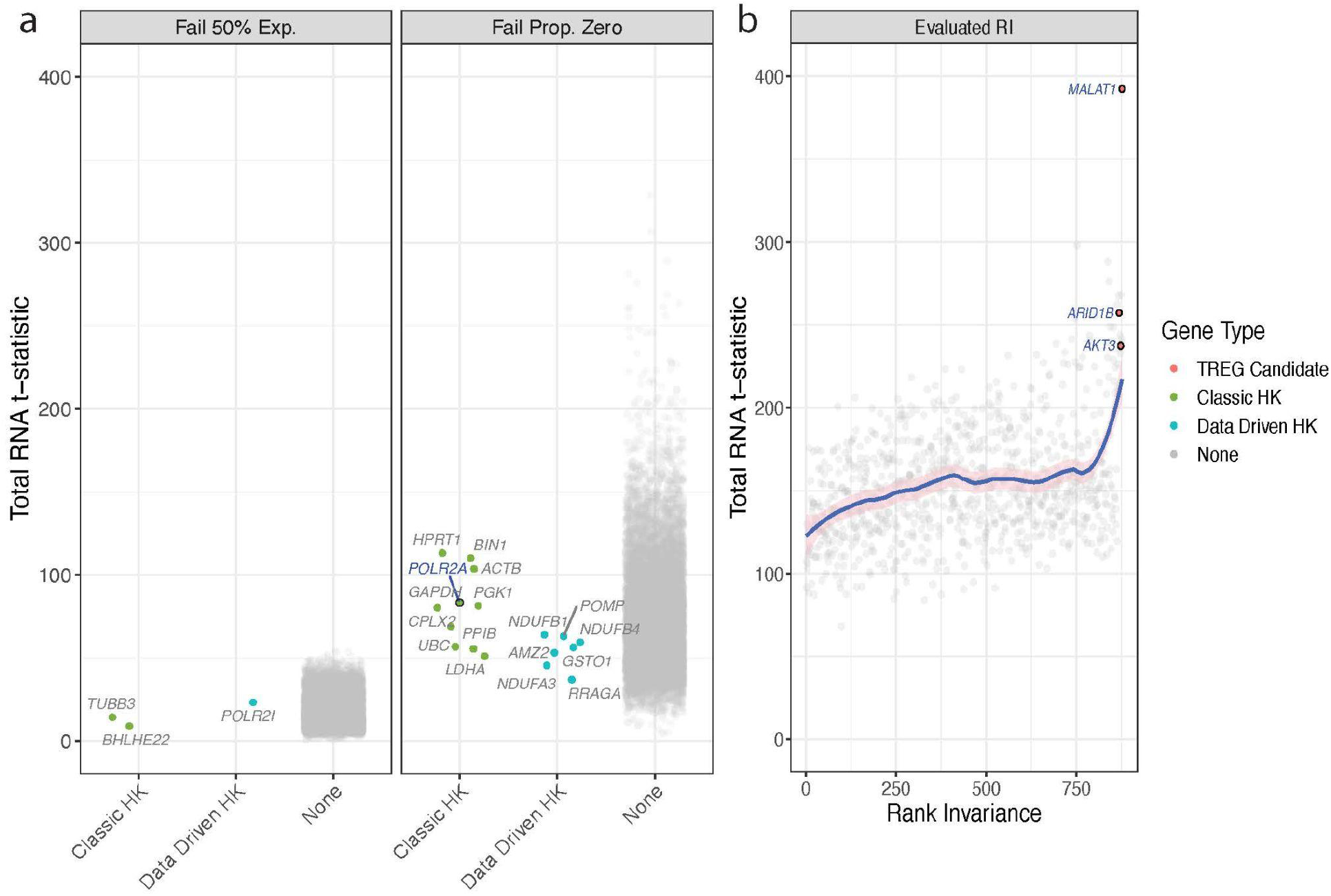
Relationship between Gene Properties and total RNA *t*-statistics. *t*-statistics are calculated for the relationship between total RNA expression (explanatory variable) and gene expression (response variable), adjusted for cell type (Methods: Total RNA linear regression) for all 23,038 genes in the reference snRNA-seq data set. **a.** The distribution of total RNA *t*-statistics for genes that failed either the top 50% filter (left panel) or the Proportion Zero filtering process (right panel). The genes are annotated by color for classic housekeeping (HK) genes [41, 42], brain data-driven housekeeping (HK) genes [6], or belonging to none of those three groups. **b.** Scatterplot of the relationship between the Rank Invariance and the total RNA *t*-statistic for the 877 genes that pass all the filtering steps. Candidate TREGs explored with RNAscope are highlighted in red and labeled. A loess regression line with span = 0.3 is shown. All three panels are mutually exclusive categories of genes based on the filtering processes. Related to **Figure 2b** and **Figure 3c-d**.

**Supplementary Figure 3:**
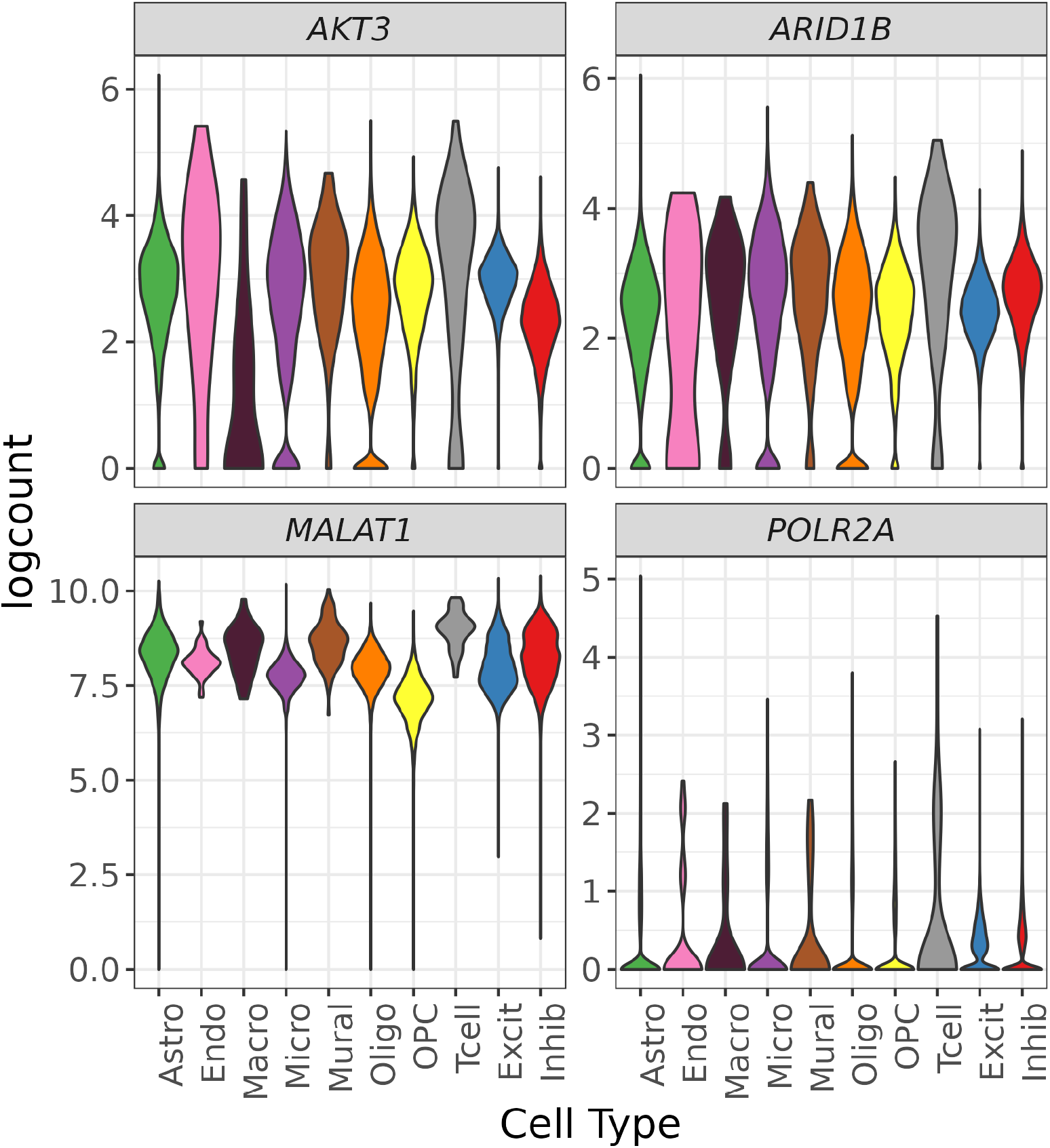
snRNA-seq expression plots. Distribution of the normalized log2-transformed expression over cell types for *AKT3, ARID1B, MALAT1*, and *POLR2A*. Candidate TREGs show less expression variability across most cell types compared to *POLR2A*. Expression Ranks (**Figure 3b**) are less variable than measured expression values since the Expression Rank takes into consideration the context of the expression levels of other genes in a given cell.

**Supplementary Figure 4:**
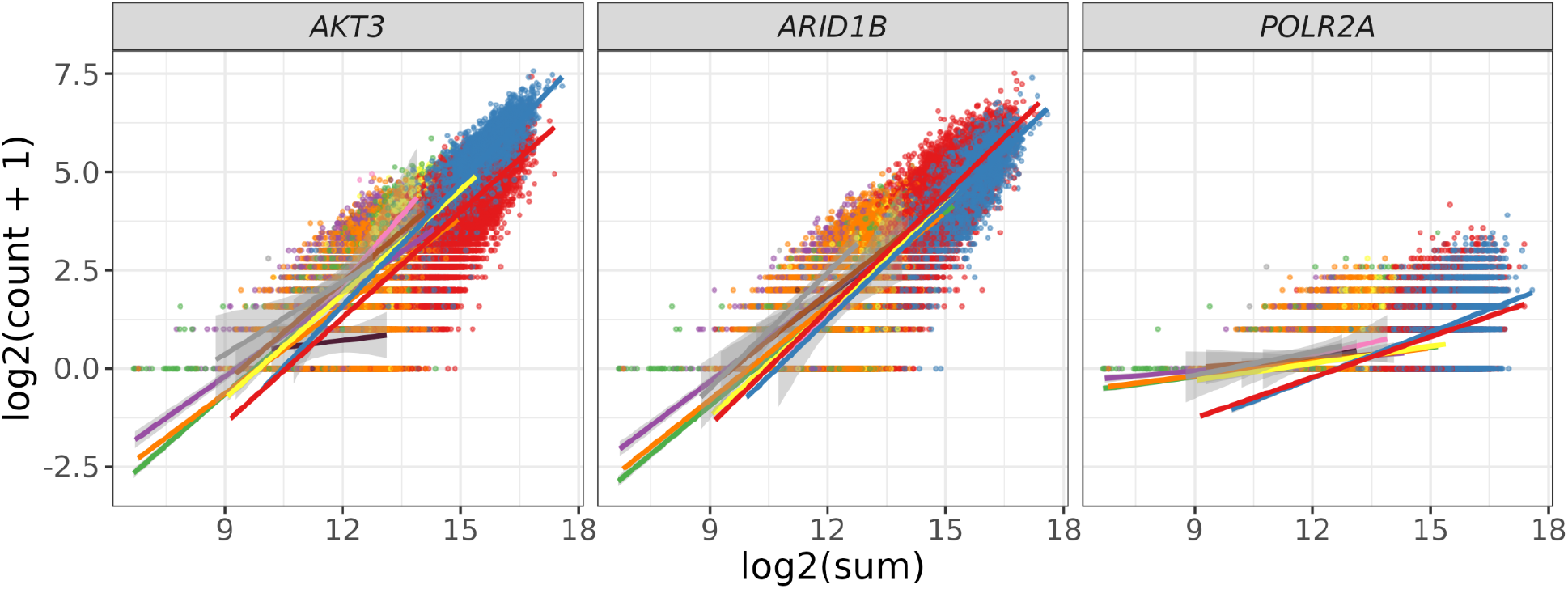
Relationship between total nuclear expression and expression of candidate TREGs. Scatter plot of the total RNA expression (estimated by the nuclei log2 sum of all counts) against the nuclei gene expression (log2 of the count plus one), and *AKT3, ARID1B, and POLR2A;* related to **Figure 3d**. The scatter plots are overlaid with the linear fit for each cell type and colored by cell type. See **Figure 3c** for *MALAT1*.

**Supplementary Figure 5:**
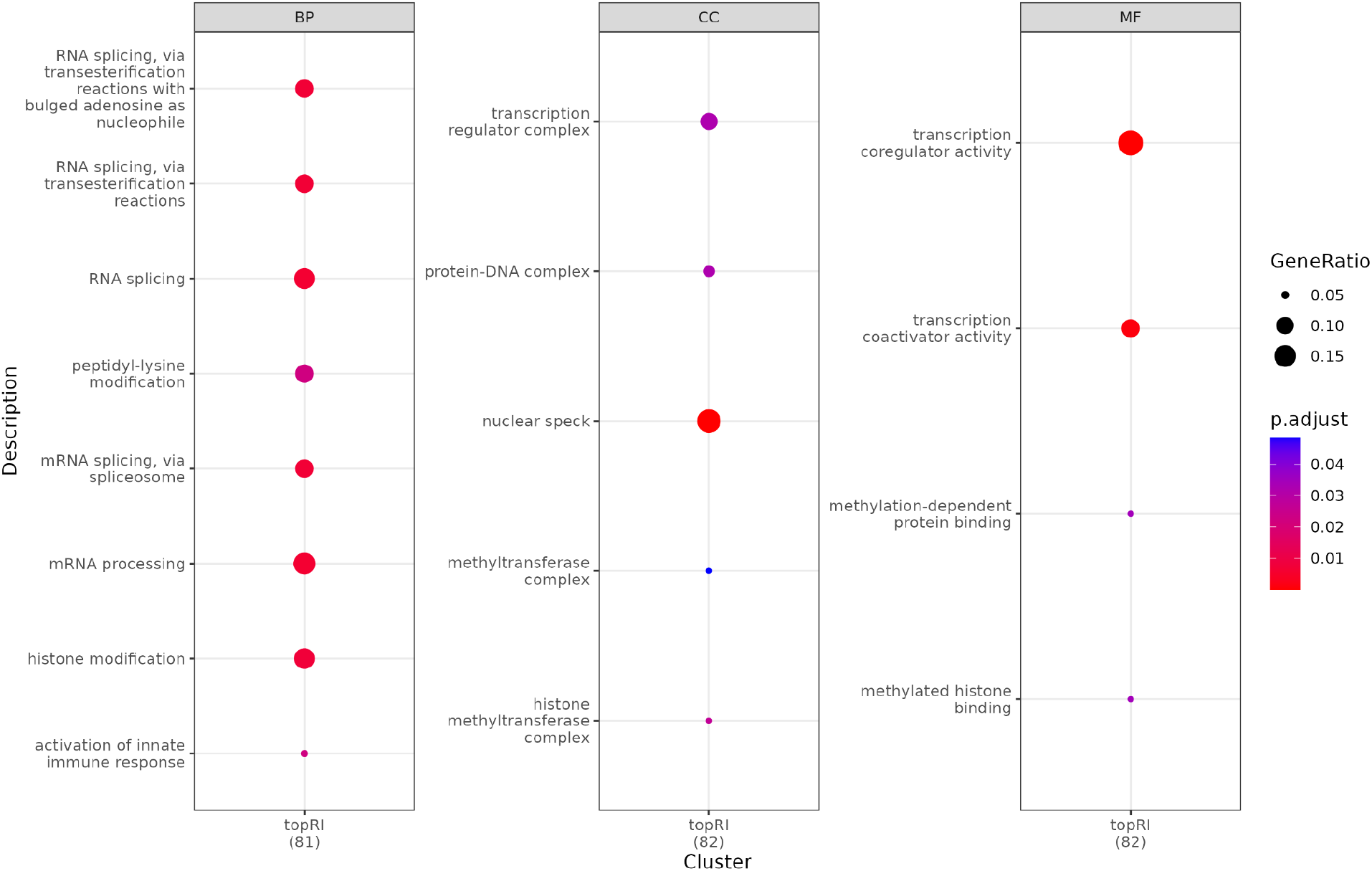
Gene ontology dotplots. Gene ontology enrichment results for the top 10% candidate TREGs found using snRNA-seq data from 5 human brain regions across ontologies for biological processes (BP), cellular components (CC), and molecular function (MF). The top 10% candidate TREGs are enriched for functions related to RNA transcription, namely RNA splicing and histone modification, as well as related cellular components and molecular functions. This finding is consistent with the TREG property that its expression is associated with transcriptional activity. Related to **Figure 3c-d**.

**Supplementary Figure 6:**
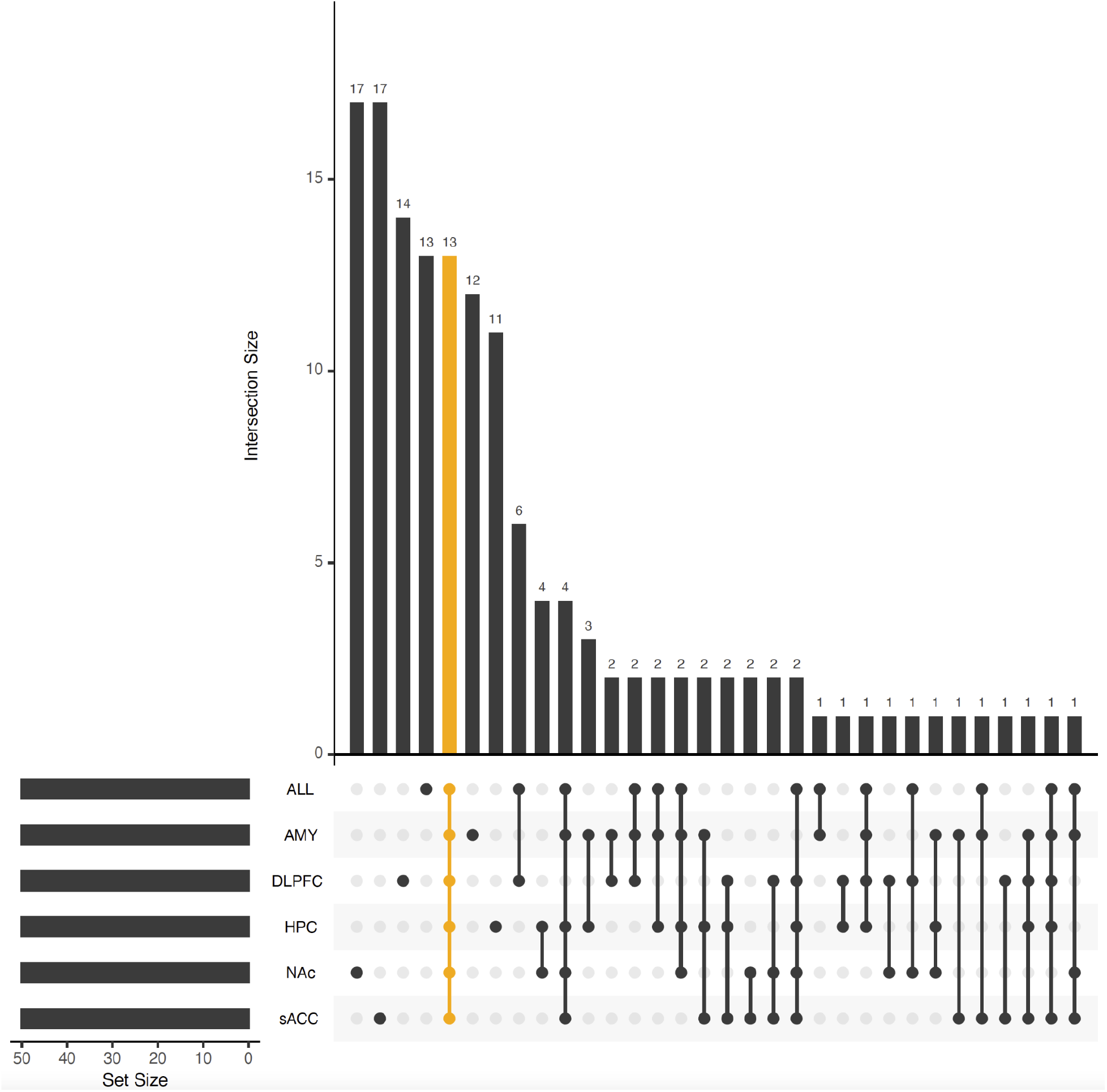
Upset plot for top 50 RI genes. The Proportion Zero filtering and Rank Invariance calculation was performed on each of five brain regions and across all brain regions. The overlap of the sets of the top 50 Rank Invariance genes from each analysis is shown in this “UpSetR” plot [40]. An “UpSetR” plot is similar to a Venn diagram as it shows bars denoting the intersection size (top) for different types of intersections (bottom right) based on sets of features (in this case gene IDs) that could have different sizes (bottom left, in this case all six sets are of equal size: 50 genes). “UpSetR” plots can include more sets than Venn diagrams. The intersections are ordered by decreasing intersection size, and here the fifth largest intersection corresponds to the intersection across all six sets (shown in orange). The genes *AKT3, ARID1B*, and *MALAT1* are part of the set of 13 genes that are present in each set of top 50 Rank Invariance genes (**Supplementary Table 2**).

**Supplementary Figure 7:**
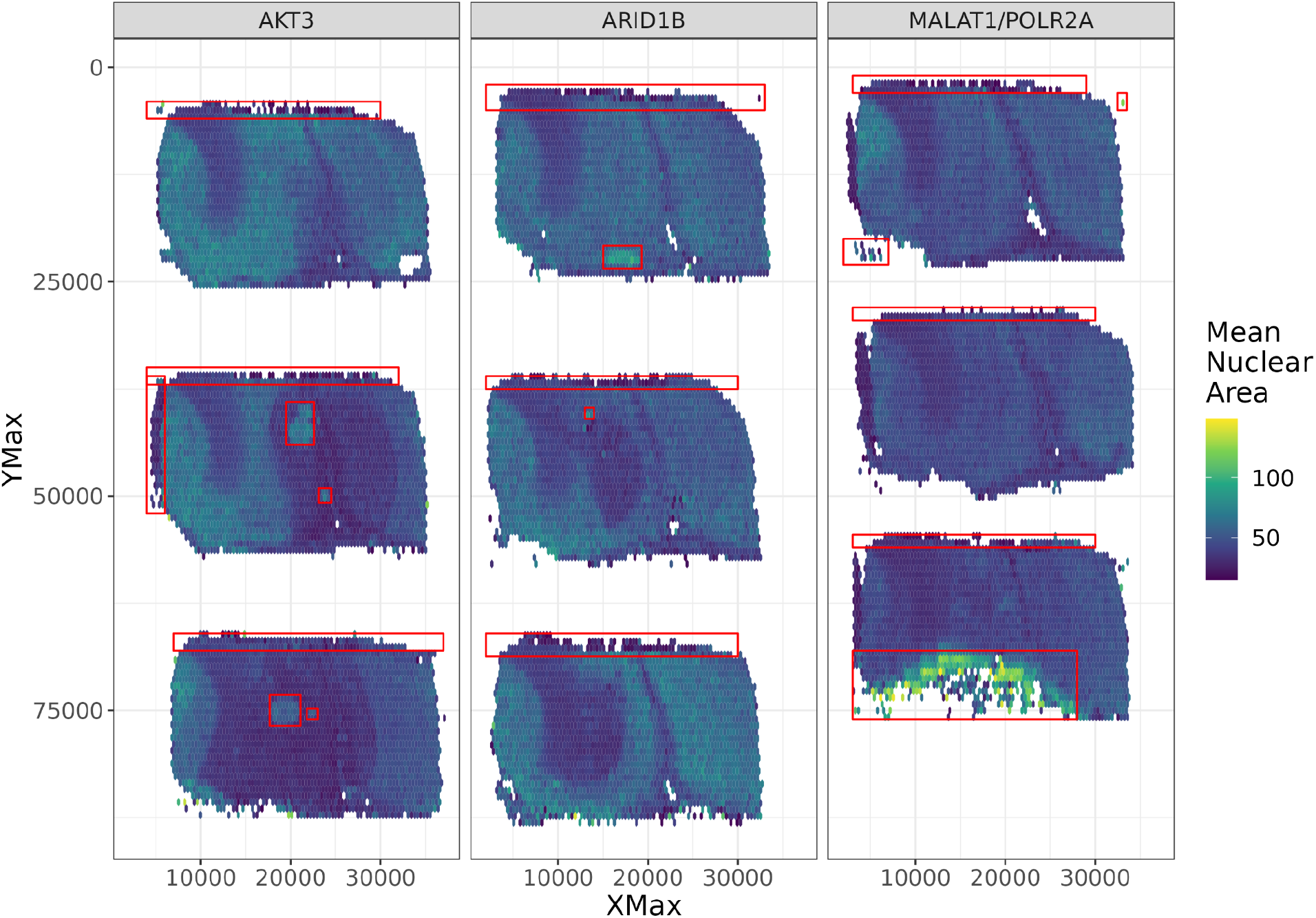
Spatial distribution of mean nuclear area across all *n* = 9 tissue sections. To visualize the nuclear area spatially, we used hex bins to compute the mean nuclear area per hex bin. The regions inside of the red boxes were flagged during quality control for unusually high nuclear size, and the enclosed nuclei were excluded from the analysis for technical artifacts due to sampling and imaging. Related to **Figure 6a**, Methods: Quality control and spatial quantification of HALO data.

**Supplementary Figure 8:**
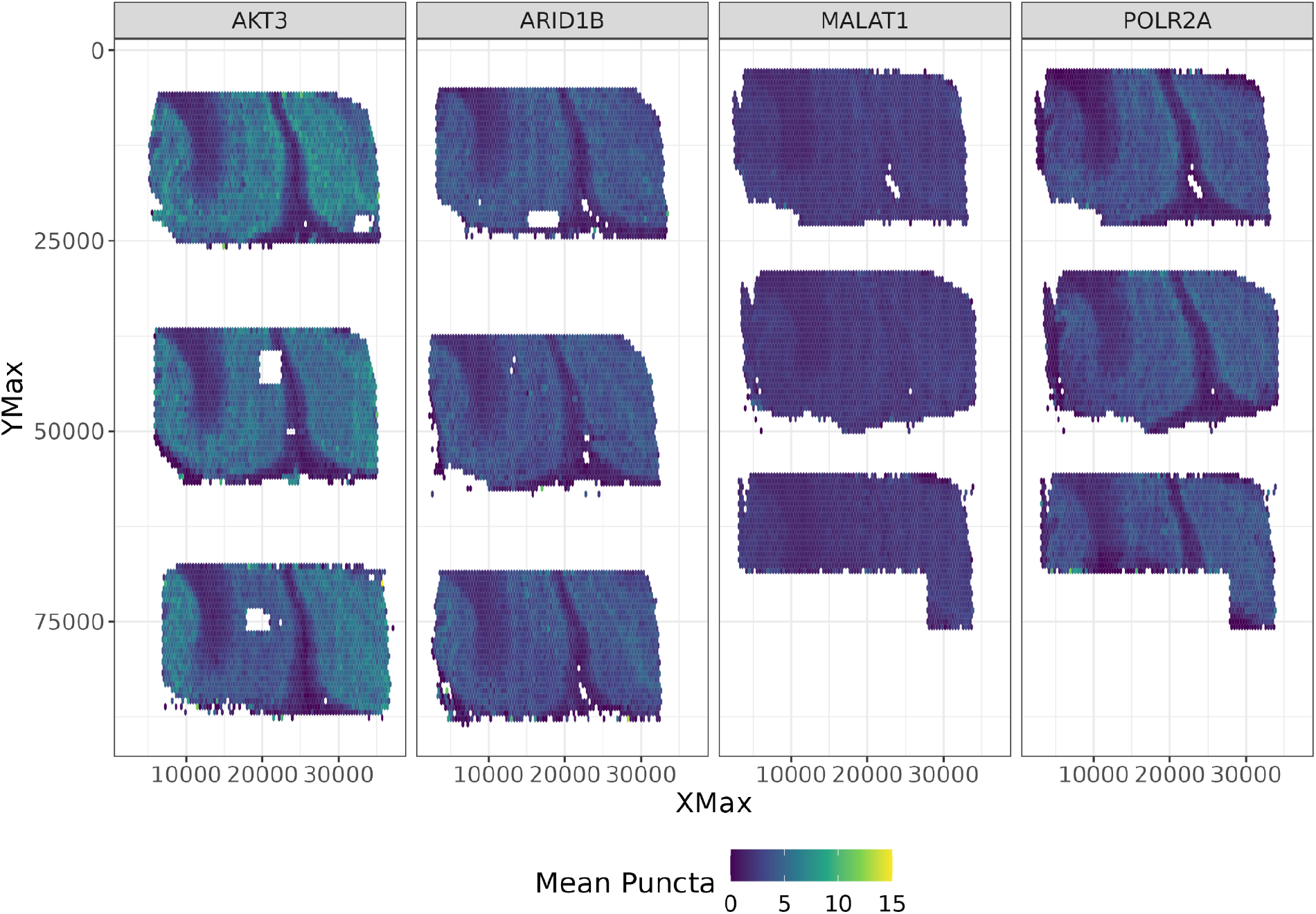
Spatial distribution of mean number of puncta across all *n* = 9 tissue sections, post quality control. Similar to **Supplementary Figure 7**, but for the mean number of puncta for each gene after performing quality control. Given that *MALAT1* and *POLR2A* were multiplexed in the same experiment on a single slide, (**Figure 4a**, **Supplementary Table 3**), their spatial shapes are identical. Related to **Figure 6b**.

**Supplementary Figure 9:**
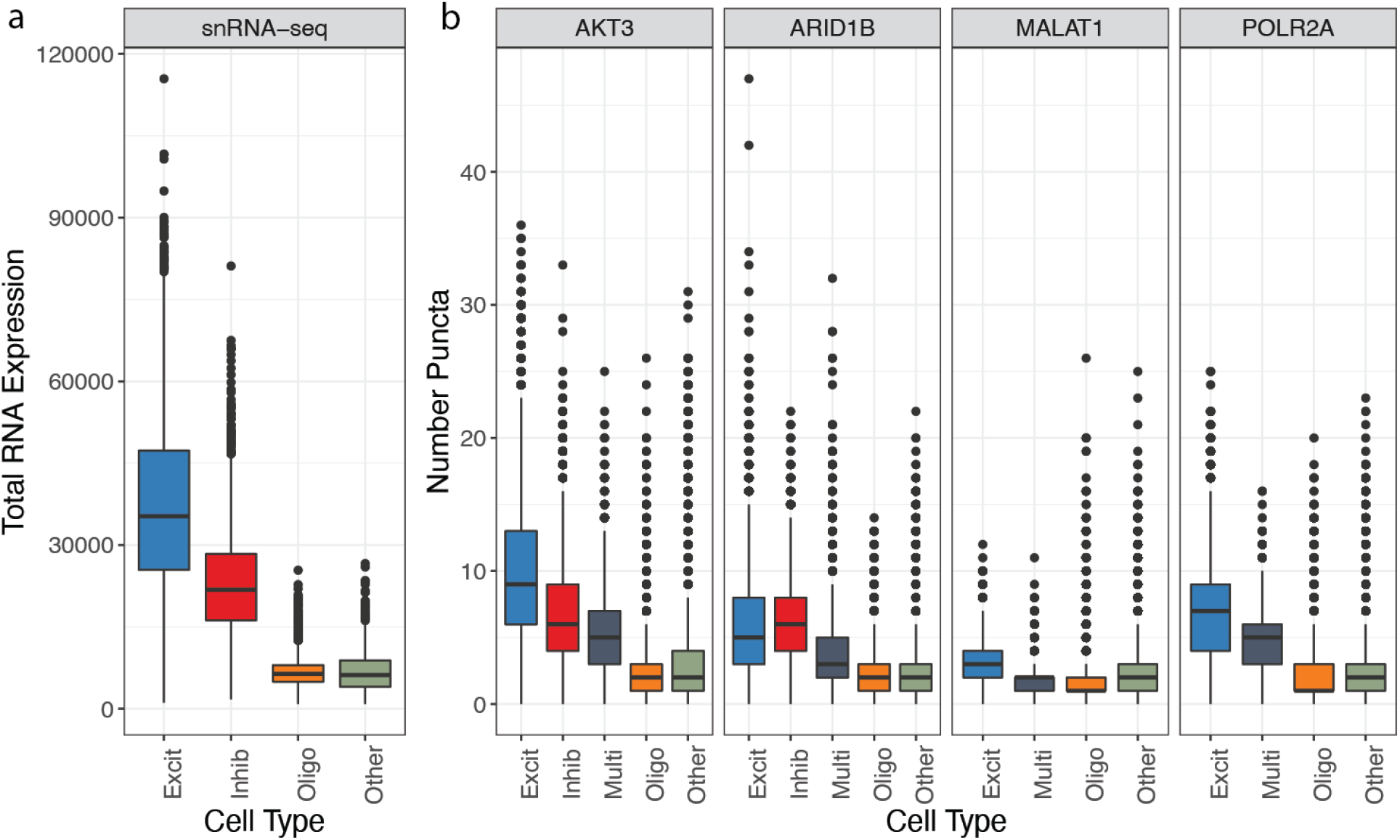
Boxplots of total nuclear RNA expression across all cell types. **a.** Total nuclear RNA expression summed from snRNA-seq data. Nuclei not classified as Excit, Inhib, or Oligo were classified as “Other’’ and likely represent other non-neuronal cell types located in the brain, such as astrocytes and microglia. **b.** Total nuclear RNA expression estimated by the number of puncta measured by RNAscope for each observed gene. Nuclei expressing more than one cell type marker (Excit, Inhib, or Oligo) are classified as “Multi” as they cannot be definitively assigned a cell type phenotype due to overlapping fluorescent signals. Related to **Figure 7** and **Table 1**.

**Supplementary Figure 10:**
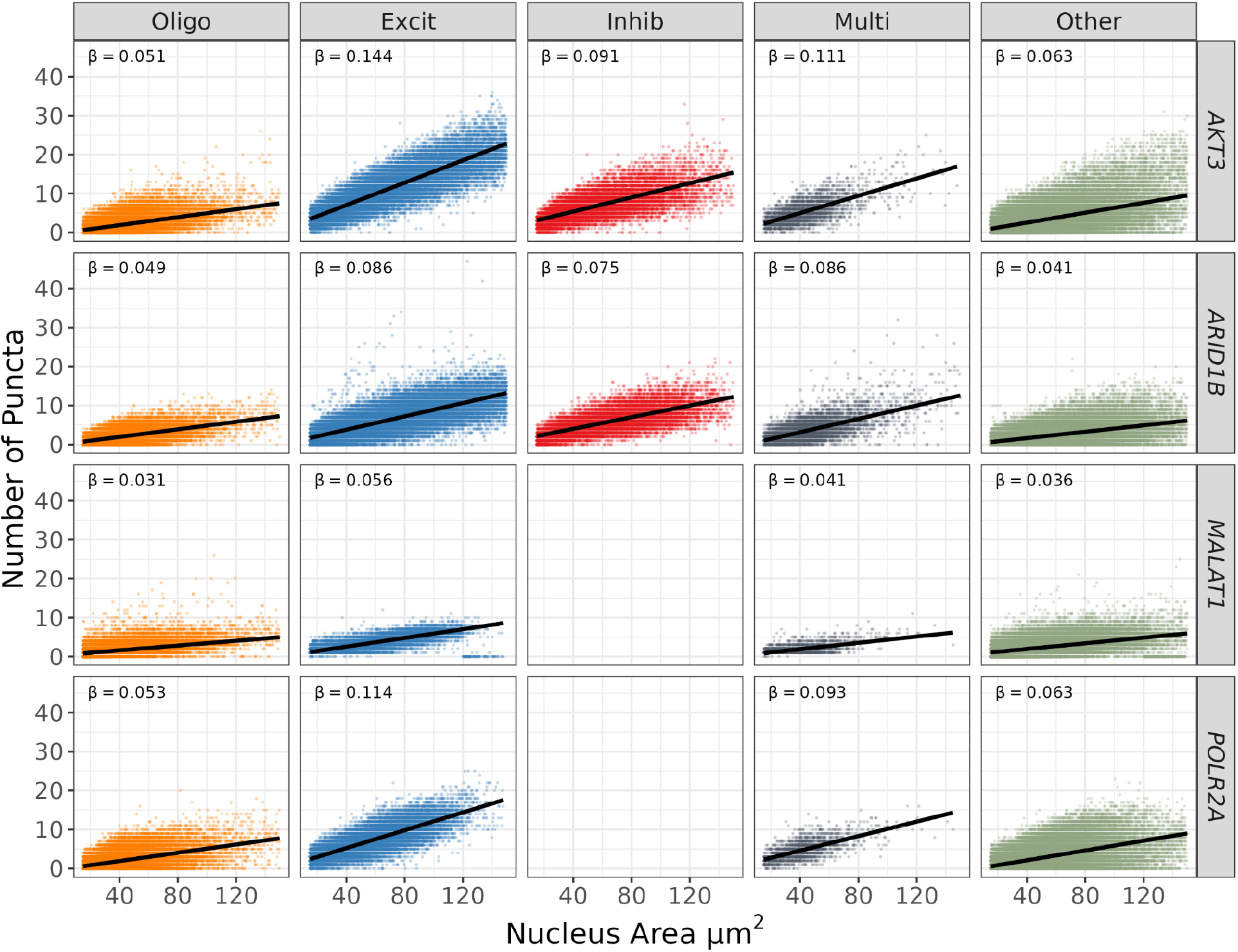
Nuclear puncta versus nuclear area across cell types. Given the paired nuclear size (area in μm^2^), RNA abundance (number of puncta), and cell type assignment data from RNAscope experiments, we can examine the relationship between puncta and area across all cell types. The relationship between nuclear area and puncta is different across cell types, for example excitatory neurons (Excit) and oligodendrocytes (Oligo) for *AKT3* have a difference in slopes of 0.093 (t=155.77, p<2*10^-16^).Cell types are the same as those from **Supplementary Figure 9** and it should be noted that *MALAT1* puncta were unreliable due to oversaturation of fluorescent signals as previously explained. Related to **Figure 7**. There is no data for inhibitory neurons for *MALAT1* and *POLR2A* as *GAD1* was omitted for this experiment due to limitations in multiplexing (**Figure 4**).

### Supplementary Tables Legends

**Supplementary Table 1**: **Numbers of different nuclei over brain regions.** Nuclei counts in the snRNA-seq dataset over the ten broad cell types and five brain regions.

**Supplementary Table 2**: **Detailed gene metrics.** Detailed gene metrics. Including the Ensembl ID, gene symbol, gene type annotation and *t*-statistic for the total RNA linear regression. For ALL regions and, and each individual region: pass the 50% expression filter (top50), the maximum Proportion Zero (max_propZero), pass the Proportion Zero filter (PropZero_filter), Rank Invariance value (rank_invar).

**Supplementary Table 3**: **RNAscope probe combinations and opal dye (fluorophore) assignments.** *GAD1* labels inhibitory neurons, *SLC17A7* labels excitatory neurons, and *MBP* labels oligodendrocytes. *AKT3, ARID1B*, and *MALAT1* are candidate TREGs while *POLR2A* is a classic HK gene.

**Supplementary Table 4: Opal dye dilutions used to fluorescently label probes in RNAscope.** Opal dyes must be diluted before application to tissue sections. Dilutions are optimized based on probe intensity (e.g. cell type marker genes with high expression require a lower concentration [higher dilution] of fluorescent Opal dyes).

**Supplementary Table 5: Scanning protocol exposure times.** The exposure times, in milliseconds, used for the Polaris scanning algorithms listed by experiment.

**Supplementary Table 6: HALO cell counts.** Columns 2-3: Numbers of cells (before and after performing quality control for poorly segmented regions) segmented by HALO software for each sample (rows) in the RNAscope experiments. Columns 4-8: The number of nuclei assigned to a given cell type category. Nuclei with more than one cell type marker (Excit, Inhib, or Oligo) are classified as “Multi” whereas those without any markers are labeled as “Other”.

## Notes

https://github.com/LieberInstitute/TREG_paper

### Competing Interest Statement

The authors have declared no competing interest.

